# Proteasomal Inhibition Triggers Viral Oncoprotein Degradation via Autophagy-Lysosomal Pathway

**DOI:** 10.1101/780171

**Authors:** Chandrima Gain, Samaresh Malik, Shaoni Bhattacharjee, Arijit Ghosh, Erle S. Robertson, Benu Brata Das, Abhik Saha

## Abstract

Epstein-Barr virus (EBV) nuclear oncoprotein EBNA3C is essential for B-cell transformation and development of several B-cell lymphomas particularly those are generated in an immuno-compromised background. EBNA3C recruits ubiquitin-proteasome machinery for deregulating multiple cellular oncoproteins and tumor suppressor proteins. Although EBNA3C is found to be ubiquitinated at its N-terminal region and interacts with 20S proteasome, the viral protein is surprisingly stable in growing B-lymphocytes. EBNA3C can also circumvent autophagy-lysosomal mediated protein degradation and subsequent antigen presentation for T-cell recognition. Recently, we have shown that EBNA3C enhances autophagy, which serve as a prerequisite for B-cell survival particularly under growth deprivation conditions. We now demonstrate that proteasomal inhibition by MG132 induces EBNA3C degradation both in EBV transformed B-lymphocytes and ectopic-expression systems. Interestingly, MG132 treatment promotes degradation of two EBNA3 family oncoproteins – EBNA3A and EBNA3C, but not the viral tumor suppressor protein EBNA3B. EBNA3C degradation induced by proteasomal inhibition is partially blocked when autophagy-lysosomal pathway is inhibited. In response to proteasomal inhibition, EBNA3C is predominantly K63-linked polyubiquitinated, colocalized with the autophagy-lsyosomal fraction in the cytoplasm and participated within p62-LC3B complex, which facilitates autophagy-mediated degradation. We further show that the degradation signal is present at the first 50 residues of the N-terminal region of EBNA3C. Proteasomal inhibition reduces the colony formation ability of this important viral oncoprotein, increases transcriptional activation of both latent and lytic gene expression and induces viral reactivation from EBV transformed B-lymphocytes. Altogether, this study offers rationale to use proteasome inhibitors as potential therapeutic strategy against multiple EBV associated B-cell lymphomas, where EBNA3C is expressed.

**Author Summary:** Epstein-Barr virus (EBV) establishes latent infection in B-lymphocytes and is associated with a number of human malignancies, both of epithelial and lymphoid origin. EBV encoded EBNA3 family of nuclear latent antigens comprising of EBNA3A, EBNA3B, and EBNA3C are unique to immunoblastic lymphomas. While EBNA3A and EBNA3C are involved in blocking many important tumor suppressive mechanisms, EBNA3B exhibits tumor suppressive functions. Although EBNA3 proteins, in particular EBNA3C, interact with and employ different protein degradation machineries to induce B-cell lymphomagenesis, these viral proteins are extremely stable in growing B-lymphocytes. To this end, we now demonstrate that proteasomal inhibition leads to specifically degradation of oncogenic EBNA3A and EBNA3C proteins, whereas EBNA3B remains unaffected. Upon proteasomal inhibition, EBNA3C degradation occurs via autophagy-lysosomal pathway, through labeling with K63-linked polyubiquitination and participating in p62-LC3B complex involved in ubiquitin-mediated autophagy substrate selection and degradation through autolysosomal process. We also demonstrate that the N-terminal domain is responsible for autophgy-lysosomal mediated degradation, while the C-terminal domain plays a crucial role in cytoplasmic localization. Fascinatingly, while proteasomal inhibition reduces EBNA3C’s oncogenic property, it induces both latent and lytic gene expressions and promotes viral reactivation from EBV transformed B-lymphocytes. This is the first report which demonstrates a viral oncoprotein degrades through autophagy-lysosomal pathway upon proteasomal inhibition. In sum, the results promise development of novel strategies specifically targeting proteolytic pathway for the treatment of EBV associated B-cell lymphomas, particularly those are generated in immunocompromised individuals.

## Introduction

Epstein-Barr virus (EBV) is a large DNA virus that belongs to the gammaherpesvirus subfamily and persistently infects majority of the worldwide human population [1,2,3]. In spite of being a ubiquitous virus, EBV is recognized as one of the most efficient transforming viruses and responsible for development of several neoplasms, mostly of B-cell origin, but also some epithelial origin such as nasopharyngeal, breast and gastric carcinomas. There are three major types of B-cell malignancies causally associated with EBV are the Burkitt’s lymphoma (BL), Hodgkin’s lymphoma (HL) and diffuse large B cell lymphomas (DLBCL) [2, 4]. EBV is also detected as the etiologic factor of majority of infectious mononucleosis (IM), a lymphoproliferative disorder attributable to a striking increase in CD8^+^ T-cell activation and subsequent expansion of B-lymphocytes upon viral infection [5]. This clinical manifestation of primary EBV infection indicates its competence not only to embark on latent infection in B-lymphocytes but also to impel B-cell proliferation *in vivo*. Moreover, *in vitro* EBV transforms nascent B-lymphocytes into continuously proliferating lymphoblastoid cell lines (LCLs) with phenotypic resemblance of activated B-blasts [6, 7]. These LCLs express similar gene sets that have been detected in EBV associated lymphoid malignancies in humans, such as immunoblasic B-cell lymphomas [1, 2]. Transformation of B-lymphocytes by EBV has been attributed to the expression of a growth stimulatory program by the virus. LCLs express all EBV latency-associated genes, including six EBV nuclear antigens (EBNAs) - EBNA1, EBNA2, EBNA3A, EBNA3B, EBNA3C and EBNALP, three latent membrane proteins (LMPs) - LMP1, LMP2A, and LMP2B), two small noncoding RNAs (EBER1 and EBER2), and microRNA transcripts from the BHRF1 and BamHI A (BART) regions [1, 2]. Out of these, five viral antigens including EBNA2, EBNA3A, EBNA3C, EBNA-LP and LMP1 act in concert to transform naïve B-lymphocytes [8,9,10,11,12]. The EBNA3 family comprising of EBNA3A, EBNA3B, and EBNA3C genes are considered to comprise a family of non-redundant EBV genes, which most likely generated from gene duplication during primate gammaherpesvirus evolution [13,14,15]. Although the EBNA3 antigens possess the similar genomic structure, they share limited amino acid sequence homology within the N-terminal region [13,14,16].

The two major proteolytic cascades that cells use for protein degradation are the ubiquitin-proteasome system (UPS) and autophagy-lysosomal pathway [17, 18]. In addition, another important family of cytosolic proteases is the caspases that cleave proteins after aspartic acid residues, also play an important role in protein degradation during apoptosis [19]. Although initially UPS and autophagy-lysosomal pathways were thought as self-governing protein degradation mechanisms, recent evidence suggests profound crosstalk between these two major protein turn-over mechanisms [18]. While the UPS typically degrades short-lived ubiquitinated proteins, autophagy-lysosomal pathway targets long-lived misfolded protein aggregates, which are produced in response to different cellular insults, such as growth factor deprivation, hypoxia and endoplasmic (ER) stress due to unfolded protein response (UPR) [20,21,22]. Autophagy is initiated by formation of a double membrane vesicle that encloses polyubiquitinated cargo protein aggregates through interacting with specific adaptor proteins, such as p62/SQSTM1. Subsequently, p62 interacts with microtubule-associated proteins 1A/1B light chain 3B (LC3B) via LC3-interacting region (LIR) motif [23, 24]. LC3B is initially synthesized in an unprocessed form, pro-LC3, which is then cleaved at the C-terminal glycine residue into cytosolic form, LC3-I. Conjugation of phosphatidyl-ethanolamine (PE) to the C-terminal residue of LC3-I generates the membrane associated form, LC3-II, providing an authentic marker for autophagosome formation [25]. Subsequently, autophagosomes fuse with the lysosome to form autolysosome, where the acidic and hydrolytic environment helps to degrade the sequestered cargo protein aggregates [25, 26].

The current study addressed the potential crosstalk among different degradation pathways with respect to EBNA3C degradation in EBV transformed B-lymphocytes. Previously it has been shown that EBNA3C can manipulate UPS either to stabilize cell oncoproteins (viz. Mdm2, Cyclin D1, c-Myc, Pim1) or enhance degradation of tumor suppressor proteins (viz. p53, p27^Kip1^, pRb, p21^CIP1^) [27,28,29,30,31,32]. In addition, EBNA3C is poly-ubiquitinated at the N-terminal region and interacts with the C8/α7 subunit of the 20S proteasome [28, 33]. While *in vitro* EBNA3C along with two other family members are all degraded in the presence of 20S proteasome, in growing LCLs, EBNA3 proteins appear to be extremely stable, with no sign of UPS-mediated degradation [33]. Moreover, EBNA3C can bypass autophagy-lysosomal mediated degradation and subsequent antigen presentation for T-cell recognition [34, 35]. Nevertheless, the underlying proteolytic mechanism that governs EBNA3C’s turn-over remains yet to be determined. Recently, we have shown that EBNA3C transcriptionally activates autophagy machinery particularly when cells are under stressed conditions [36]. This autophagy activation appears to be essential for EBNA3C mediated latently infected B-cell survival. Moreover, EBNA3C expression in B-lymphocytes leads to accumulation of K63-linked poly-ubiquitinated protein aggregates directed for autophagy-lysosomal mediated degradation [36]. Herein we sought to investigate how EBNA3C protein degradation is regulated and the potential for targeting degradative pathways as a treatment strategy for various EBV associated B-cell lymphomas where EBNA3C is expressed.

## Results

### Proteasomal inhibition facilitates degradation of EBV essential nuclear antigen EBNA3C

Proteasomal inhibition leads to degradation of many important cell oncoproteins through autophagy-lysosomal pathway [37, 38]. For example, expression of oncogenic mutant p53 is significantly suppressed in cancer cells after treatment with proteasome inhibitors [37]. The prominent interaction of EBV essential antigen EBNA3C with the ubiquitin-proteasome system (UPS) [30, 33] and autophagy-lysosomal pathway [36] led us to investigate the mechanism that governs the extraordinary stability of this viral oncoprotein in growing EBV transformed B-lymphocytes.

In order to determine the half-life of EBNA3C protein affected by UPS and autophagy-lysosomal pathway, EBV transformed lymphoblastoid cell line (LCL#1) was treated with protein synthesis inhibitor, cyclohexamide (CHX) for 24 h in the absence (DMSO control) or in the presence of proteasome (MG132, 1 µM) or autophagy-lysosomal (Chloroquine, CQ, 50 µM) inhibitors (Fig. 1A). Beyond 24 h, the experiments became unreliable due to significant cell death induced by the treatment of different drugs. Samples were analyzed by western blotting (WB) of total protein extracts solubilized in lysis buffer and probing with antibodies directed against EBNA3C and GAPDH as loading control. Unexpectedly, the results demonstrated that instead of increasing EBNA3C stability, MG132 treatment resulted in a decrease of EBNA3C stability (half-life: ∼12 h) in comparison to the DMSO treated cells (half-life: >24 h), whereas CQ treatment enhanced the overall EBNA3C protein stability (Fig. 1A). To rule out the possible effect of MG132 in transcriptional regulation, a similar experimental set up was carried out using EBNA3C stably expressing BJAB cells (BJAB-E3C#7) (Fig. 1B). As before, a parallel result was obtained (Fig. 1B), indicating that MG132 mediated EBNA3C’s degradation is likely controlled at the protein level.

**Figure 1:**
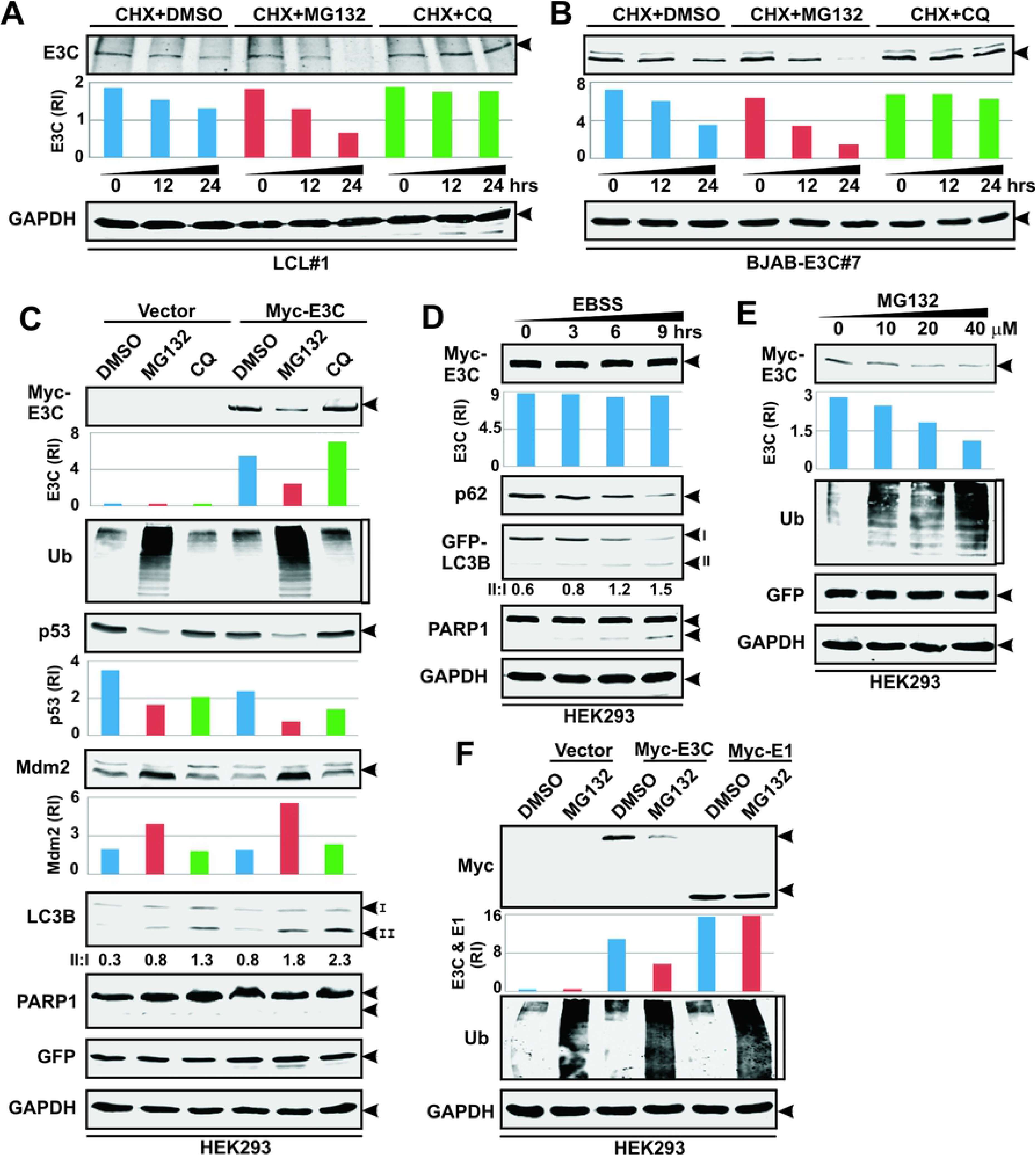
Proteasome inhibition leads to degradation of EBV essential antigen EBNA3C. (A) ∼10 x 10^6^ LCL#1 and (B) BJAB stably expressing EBNA3C (BJAB-E3C#7) cells were treated with protein synthesis inhibitor, cyclohexamide (CHX) for 24 h in the absence (DMSO control) or in the presence of 1 µM proteasome inhibitor MG132 or 50 µM autophagy-lysosomal inhibitor Chloroquine (CQ). (C) ∼10 x 10^6^ HEK293 cells transiently transfected with empty vector (pA3M) or pA3M-EBNA3C expressing myc-tagged EBNA3C. 36 h post-transfection cells were either left untreated (DMSO control) or treated with 20 µM MG132 or 50 µM CQ for another 4 h before harvesting. HEK293 cells transfected with myc-tagged EBNA3C were incubated with (D) EBSS to induce autophagy for 9 h or (E) increasing concentrations of MG132 (0-40 µM) for 4 h. (F) HEK293 cells transiently transfected with empty vector or expression plasmids for myc-tagged EBNA3C or EBNA1, were either left untreated or treated with 20 µM MG132 for additional 4 h. (A-F) In each experiment, cells were harvested after drug treatment, washed with 1 x PBS, lysed in RIPA and fractionated using appropriate SDS-PAGE. For transient transfection studies, cells were additionally transfected with GFP expression vector to monitor the transfection efficiency. Western blots were performed with indicated antibodies. GAPDH blot was performed as loading control. The relative intensities (RI) of protein bands shown as bar diagrams were quantified using the software provided by Odyssey CLx Imaging System. Representative gel pictures are shown of at least two independent experiments.

Autophagy activation has been viewed as a compensatory protein degradation mechanism when proteasome function is inhibited [18, 39]. In agreement to this, our results also demonstrated that MG132 treatment significantly increased LC3II conversion along with accumulation of poly-ubiquitinated proteins in HEK293 cells transiently transfected with either vector control or myc-tagged EBNA3C (Fig. 1C). Autophagy activation upon MG132 treatment appears to be directly correlated with EBNA3C’s degradation (Fig. 1C). Upon normalization with GAPDH as loading control and GFP as transfection efficiency control, protein quantification revealed about 2 fold reduction in EBNA3C amount in MG132 treated cells as compared to DMSO treated cells, whereas CQ treatment caused no change or slight enhancement of EBNA3C amount as similar to the B-cells (Fig. 1C). Earlier it has been demonstrated that MG132 treatment induces autophagy-lysosomal mediated degradation of mutant p53 in breast cancer cell lines [37]. Although p53 is wild-type in HEK293 cells [40], a similar degradation pattern was observed in our experiment (Fig. 1C). However, Mdm2 expression was accumulated in MG132 treated cells (Fig. 1C). Interestingly, no sign of EBNA3C’s degradation was observed when autophagy was triggered in absence of growth promoting signals (Earle’s Balanced Salt Solution, EBSS) for 9 h (Fig. 1D). In contrast, a dose dependent degradative pattern of EBNA3C was observed with increasing concentrations of MG132 (0-40 μM) treated for 4 h (Fig. 1E).

Previously, it has been shown that an internal glycine-alanine repeat domain of EBNA1 plays a critical role in inhibiting UPS mediated degradation as well as acts as a cis-inhibitor for MHC class I-restricted presentation through circumventing autophagy-lysosomal degradation mechanism [34,41,42]. In contrast to EBNA3C, no sign of degradation was observed for EBNA1 in response to MG132 treatment of transiently transfected HEK293 cells with (Fig. 1F). Overall, the data suggest that proteasomal inhibition by MG132 enhances EBNA3C’s degradation and autophagy-lysosomal pathway might play an important role in regulating EBNA3C stability.

### Proteasomal inhibition leads to degradation of latent oncoproteins - EBNA3A and EBNA3C, but not EBNA3B

EBNA3 family members comprising EBNA3A, EBNA3B and EBNA3C share approximately 30% sequence homology near the N-terminal region [16]. All three EBNA3 proteins interact with C8 (α7) subunit of the 20S proteasome and are efficiently degraded *in vitro* in the presence of purified 20S proteasome [33]. However, *in vivo* in proliferating LCLs no sign of proteolytic degradation was observed for these proteins [33]. Moreover, EBNA3 proteins exert extensive cooperative functions and sometimes opposing activities [43, 44]. While EBNA3A and EBNA3C are considered as viral oncoproteins by largely collaborating in growth promoting functions, EBNA3B exhibits tumor suppressive functions [8,9,45]. It was therefore important to determine whether EBNA3A and/or EBNA3B proteins were also similarly regulated in LCLs when proteasome was inhibited.

To this end, two LCLs – LCL#1 and LCL#89 were either left untreated (DMSO control) or treated with 1 µM MG132 and subjected to western blot analyses (Fig. 2A). Since cell viability experiment demonstrated that 1 µM MG132 treatment for 12 h did not induce significant cell mortality (∼10%) in LCLs (data not shown), unless and otherwise stated, henceforth all the experiments were carried out at this dose. The results demonstrated that MG132 treatment resulted in a prominent decrease in expressions of both EBNA3A and EBNA3C viral oncoproteins, whereas unexpectedly no sign of degradation was observed for the third member of EBNA3 family proteins - EBNA3B, a non-essential viral latent antigen during B-cell transformation, in both LCLs (Fig. 2A). A similar degradation pattern of both EBNA3A and EBNA3C was also found in BJAB cells stably expressing these viral oncoproteins (Fig. S1A). Studies suggested that EBV encoded latent transmembrane oncoprotein LMP1 is a short-lived protein and unlike other cell plasma membrane substrates, it is degraded through ubiquitin mediated proteasomal pathway and not by autophagy-lysosomal pathway. In agreement to this, our results also revealed that proteasome inhibition by MG132 considerably increased LMP1 expression as compared to the DMSO treated cells (Fig. 2A). Effect of MG132 treatment was assessed by overall accumulation of polyubiquitinated proteins, induction of apoptosis as indicated by increased PARP1 cleavage and autophagy activation as shown by increased ratio of LC3II/I (Fig. 2A). Interestingly, as similar to HEK293 experiments, MG132 treatment resulted in pronounced p53 degradation in both LCLs and BJAB cells stably expressing either EBNA3A or EBNA3C (Fig. 2A and S1). However, at this point we are not sure whether these LCLs (donor specific) express mutated or wild-type p53, since only mutated p53 was shown to be degraded in the presence of proteasomal inhibitors.

**Figure 2:**
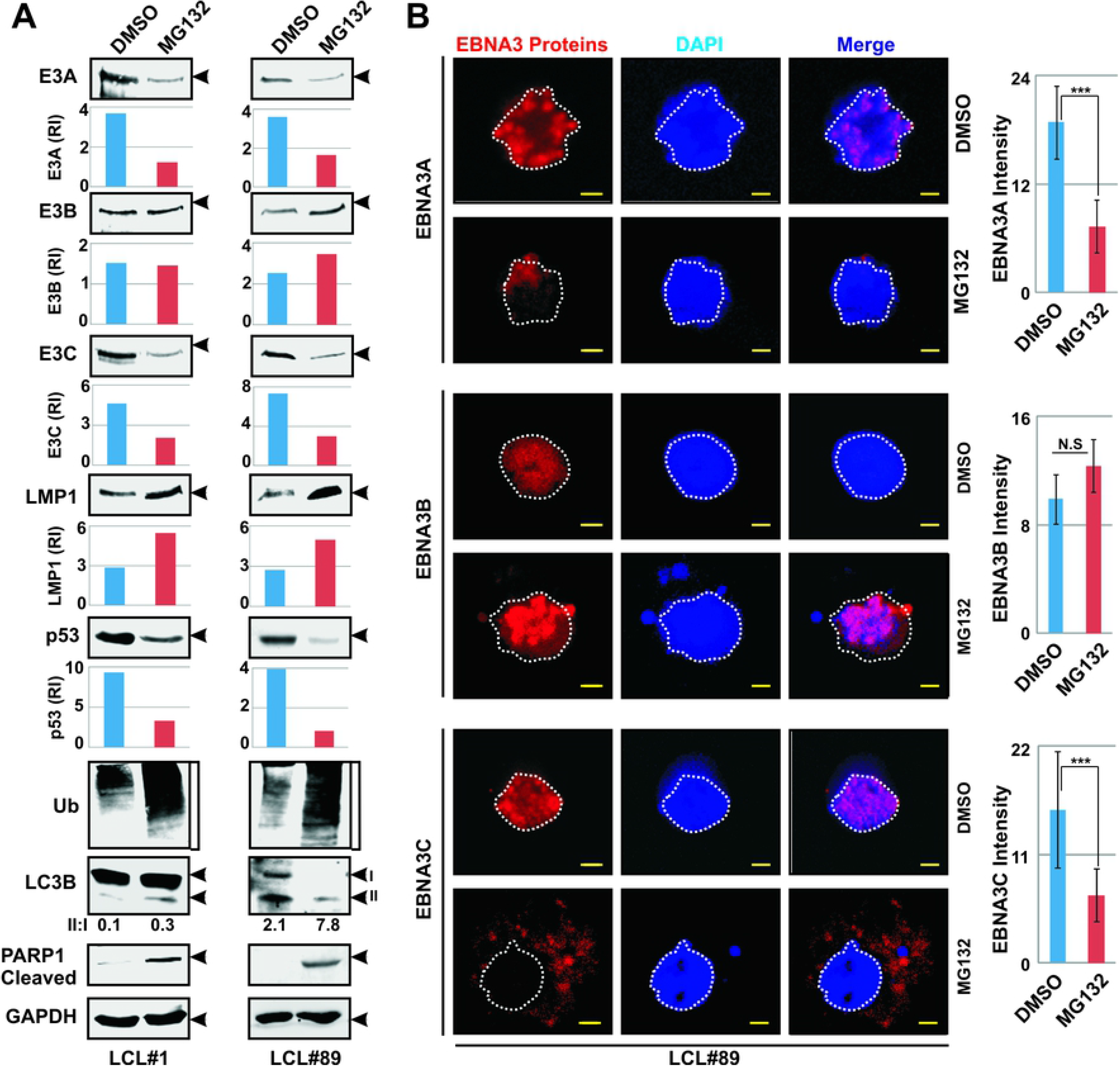
Proteasomal inhibition results in degradation of viral oncoproteins - EBNA3A and EBNA3C, but not EBNA3B in LCLs. (A) ∼10 x 10^6^ LCLs – both LCL#1 and LCL#89 were either left untreated (DMSO control) or treated with 1 µM MG132. 12 h post-incubation, cells were harvested, washed with 1 x PBS, lysed in RIPA buffer and subjected for western blot analyses with indicated antibodies. GAPDH blot was used as loading control. The relative intensities (RI) of protein bands shown as bar diagrams were quantitated using the software provided by Odyssey CLx Imaging System. Representative gel pictures are shown of at least two independent experiments. (B) For confocal assays, ∼5 x 10^4^ LCLs (LCL#89) either left untreated (DMSO control) or treated with 1 µM MG132 for 12 h, were fixed with 4% paraformaldehyde. Immuno-staining was performed using primary antibodies against viral proteins - EBNA3A, EBNA3B and EBNA3C followed by Alexa Fluor conjugated secondary antibodies for visualization in a Leica DMi8 Confocal Laser Scanning Microscope. All panels are representative pictures of two independent experiments. The bar diagram represents the mean value of staining intensities of EBNA3 proteins from at least 10 cells of 3 different fields. *** indicates P < 0.05. Nuclei were counterstained using DAPI (4’,6’-diamidino-2-phenylindole) before mounting the cells.

To corroborate proteasome inhibition mediated degradation of EBNA3 proteins, an immunofluorescence study was performed in LCL#89 without (DMSO) or with 1 µM MG132 treatment for 12 h (Fig. 2B). As similar to the western blot analyses, the immunofluorescence results also demonstrated that MG132 treatment resulted in a decrease of staining intensities (∼50%) for both EBNA3A and EBNA3C, but not EBNA3B (Fig. 2B). To confirm or rule out the involvement of proteasome inhibition in EBNA3 protein degradation, another proteasome inhibitor bortezomib, a FDA approved drug against multiple myeloma [46], was also tested. Using a similar experimental set up, the results demonstrated that alike MG132, bortezomib also induced an effective degradation phenomenon of both EBNA3A and EBNA3C proteins (Fig. S1B). Taken together, the results suggest that, impairment of proteasomal degradation machinery leads to specific degradation of EBV oncoproteins EBNA3A and EBNA3C and not the viral tumor suppressor protein EBNA3B through employing other proteolytic mechanism(s).

### Proteasomal inhibition induces both viral and autophagy gene transcriptions

A number of potential therapeutic strategies have been described with transcriptional activation of EBV lytic genes using multiple chemotherapeutic agents such as HDAC inhibitor (sodium butyrate; NaBu) in combination with a protein kinase C activator (12-O-tetradecanoylphorbol-13-acetate; TPA) [47,48,49]. Additionally, proteasome inhibitor, bortezomib treatment can also enhance EBV lytic gene transcription where CCAAT/enhancer-binding protein β (C/EBPβ) plays an important role in transactivating BZLF1 through binding to its promoter region [50]. However, the effect of proteasomal inhibition on other viral genes – both lytic and latent was not explored. To evaluate the effect of proteasomal inhibition at the transcriptional level of viral genes, quantitative reverse transcription–polymerase chain reaction (qRT–PCR) assays were performed in MG132-treated and control LCLs (Fig. 3A and 3C). The results demonstrated that MG132 treatment robustly enhanced both lytic and latent genes transcription including EBNA3A and EBNA3C, in contrast to its effect at their protein levels (Fig. 3A and 3C). Transcriptional activation of lytic genes led us to further investigate whether MG132 treatment could also induce EBV lytic cycle activation. In agreement to the previously demonstrated experiment with bortezomib, our results also showed a similar trend of EBV lytic cycle activation in both MG132 and NaBu/TPA treated LCLs compared to control cells (Fig. 3B and 3D).

**Figure 3:**
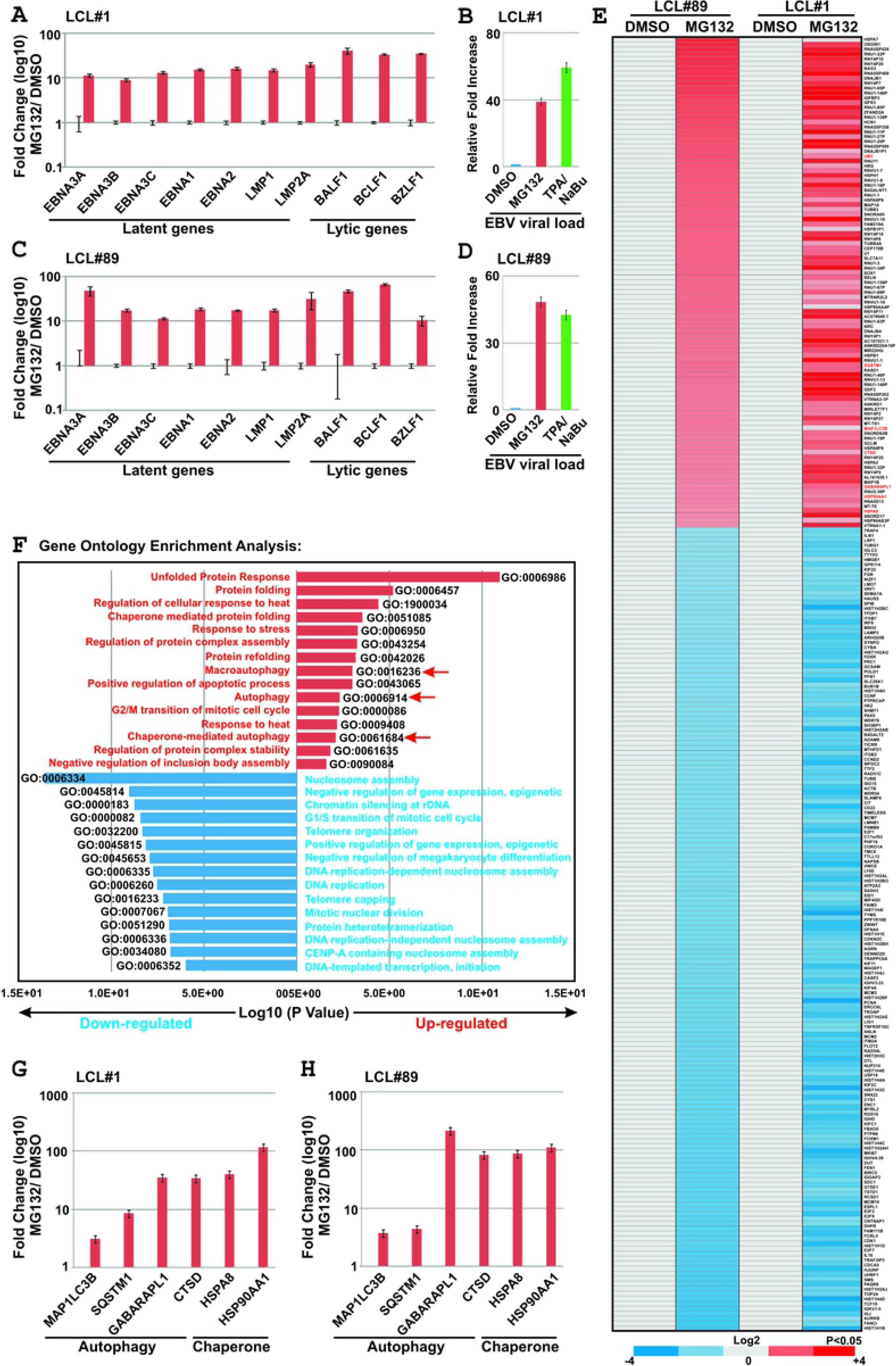
Proteasomal inhibition induces both viral and autophagy gene transcriptions. (A-D) ∼10 x 10^6^ two LCL clones – LCL#1 and LCL#89 were either left untreated (DMSO control) or treated with 1 µM MG132. 12 h post-treatment cells were harvested for (A and C) total RNA or (B and D) genomic DNA isolation as described in the “Materials and Methods” section. (B and D) LCLs were treated with 3 mM sodium butyrate (NaBu) in combination with 20 ng/ml 12-O-tetradecanoylphorbol-13-acetate (TPA) for 24 h to induce viral lytic cycle as positive control. (A and C) Total RNA was subjected to cDNA preparation followed by quantitative real-time PCR (qPCR) analyses for the selected viral genes. The relative changes in transcripts (log10) using the 2^−ΔΔCt^ method are represented as bar diagrams in comparison to DMSO control using GAPDH, B2M and RPLPO as housekeeping genes. Two independent experiments were carried out in similar settings and results represent as an average value for each transcript. (B and D) qPCR was performed for the detection of EBV DNA (BamHW fragment) using the genomic DNA isolated from each sample. The average fold increase of two independent experiments represented as bar diagrams was calculated in comparison to DMSO control using the 2^−ΔΔCt^ method taking GAPDH as housekeeping genes. (E) The cDNA samples similarly prepared in (B and D) were subjected to whole transcriptome analyses (RNA-Seq) using Ion S5^TM^ XL System as described in the “Materials and Methods” section. RNASeqAnalysis plugin (v5.2.0.5) was utilized to perform the analysis after align with human genome (hg19) and produce gene counts for all the samples. Differential gene expressions were performed based on p-value as <=0.05 and log_2_ Fold Change as 2 and above (upregulated, red) and −2 and below (downregulated, blue). (F) Differentially expressed gene sets were uploaded on DAVID v6.8 webserver for functional analysis. Gene Ontology (GO) was selected from the hits table for DAVID clustering. The bar diagrams (upregulated: red; downregulated: blue) represent top 15 most significantly affected pathways. (G) qPCR analyses of the selected cellular genes as described in (A and C).

To further investigate the global effect of proteasomal inhibition at transcriptional level, RNA-seq experiment was conducted and analyzed the transcript levels of genes that are deregulated in LCLs upon MG132 treatment (Fig. 3E-F and Table S1). The results demonstrated that upon MG132 treatment there was a significant increase in the transcripts levels of genes involved in either autophagy activation (p62/*SQSTM1*, *MAP1LC3B*, *GABARAPL1*, *CTSD*, *UBC*), or protein folding and regulation of cellular response to various cellular insults such as unfolded protein response (UPR) and chaperone mediated autophagy (*HSPH1*, *HSP90AA1*, *HSPA2*, *HSPB1*, *DNAJB1*, *DNAJB4*, *SERPINH1*, *HSPA8*, *HSP90AA4P*, *HSP90AB3P*, *BAG3*) as compared to the control LCLs (Fig. 3E-F). On the other hand, proteasomal inhibition caused a drastic reduction of mRNA levels of genes particularly involved in DNA replication and nucleosome assembly (Fig. 3E-F). The RNA-seq data for autophagy (p62/*SQSTM1*, *MAP1LC3B*, *GABARAPL1*, and *CTSD*) and chaperone (*HSP90AA1* and *HSPA8*) gene transcriptions were further validated by qRT-PCR assays in both LCLs (Fig. 3G-H).

### Autophagy stimulated by proteasomal inhibition is required for the degradation of EBNA3C

Autophagy activation as confirmed by both western blot and RNA-Seq analyses upon MG132 treatment, prompted us to further investigate that whether autophagy-lysosomal pathway is responsible for EBNA3C degradation when proteasome is inhibited. To this end autophagy pathway was blocked either by chloroquine (CQ; 50 µM) treatment or Beclin 1 knockdown in HEK293 cells (Fig. 4A and B, respectively). CQ increases the lysosomal pH and thereby blocks protein degradation. Beclin 1 is one of the initial components that initiate autophagosome formation. HEK293 cells were stably knockdown for Beclin1 by transfecting a lentiviral sh-Beclin1 expressing construct under doxycycline inducible promoter (Fig. 4B). The results demonstrated that MG132 induced EBNA3C degradation was partially inhibited by CQ treatment and beclin 1 knockdown condition (Fig. 4A and B, respectively). A similar degradation pattern of p53 in response to proteasomal inhibition and possible involvement of autophagy pathway was also validated (Fig. 4A). Unlike EBNA3C, MG132 induced p53 (endogenous) degradation was fully restored when cells were co-treated with CQ (Fig. 4A). Both LC3B and p62 were used to evaluate the autophagy level and as expected, LC3B-II/I ratios were increased following MG132 treatment alone and with CQ, concomitantly with the EBNA3C expression levels (Fig. 4A-B). Accumulation of poly-ubiquitinated proteins were also detected in accordance with MG132 treatment (Fig. 4A-B).

**Figure 4:**
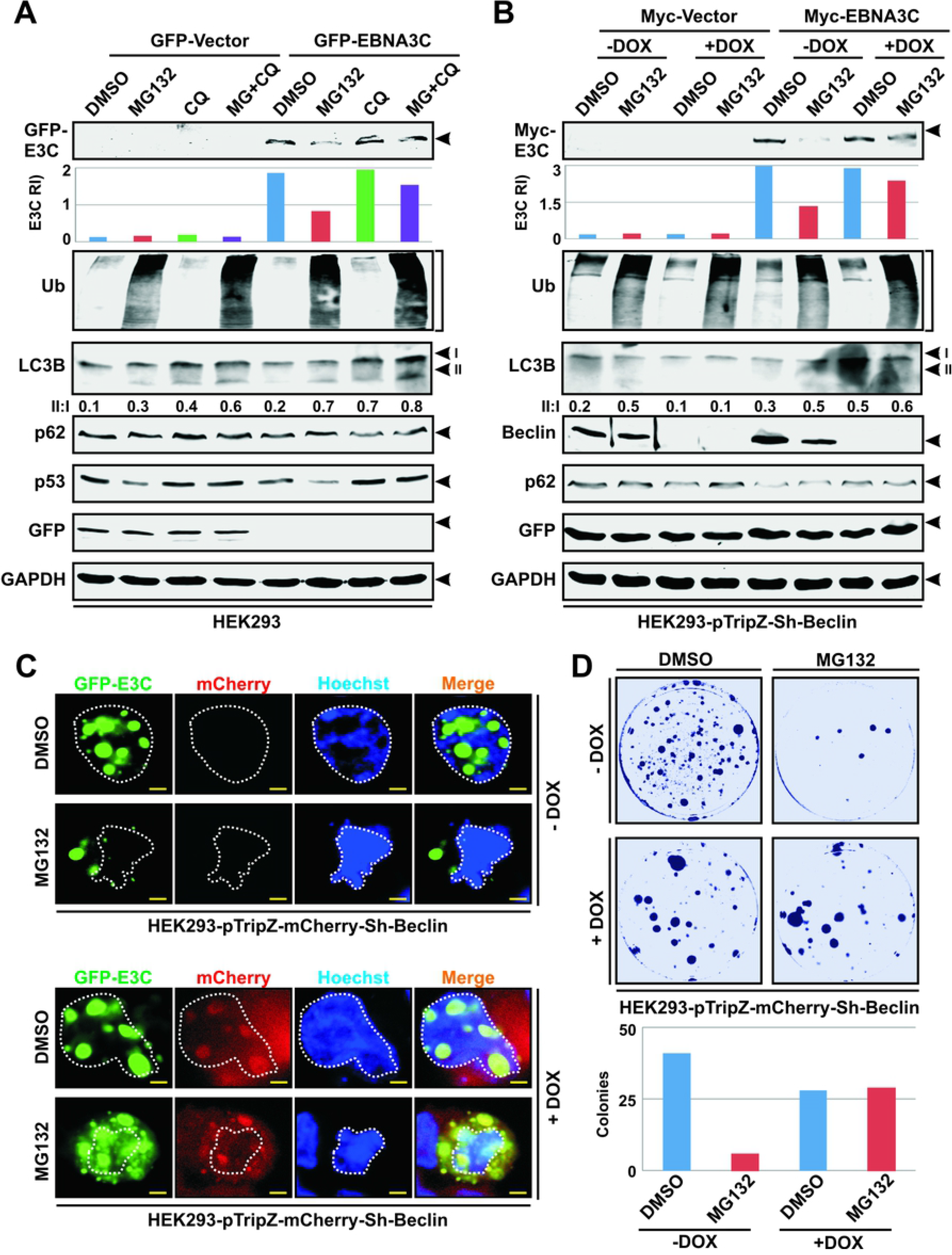
EBNA3C is degraded through autophagy-lysosomal pathway when proteasome is inhibited. (A) ∼10 x 10^6^ HEK293 cells transfected with either empty vector (pEGFP-C1) or GFP-tagged EBNA3C expression plasmid, either left untreated (DMSO control), or treated with 20 µM MG132 alone or MG132 plus 50 µM chloroquine (CQ). (B) HEK293 cells stably transfected with pTripz-mCherry-Sh-Beclin1 construct expressing sh-Beclin 1 under doxycycline (Dox) inducible promoter were further transfected either empty vector (pA3M) or myc-tagged EBNA3C expressing construct. 36 h post-transfection cells were either left untreated or treated with 20 µM MG132. (A-B) 4 h post-treatment cells were harvested, washed with 1 x PBS, lysed in RIPA buffer and subjected for western blot analyses for the indicated antibodies. (C) ∼5 x 10^4^ HEK293 cells stably expressing sh-Beclin 1 with or without doxycycline treatment were transfected with GFP-tagged EBNA3C expressing plasmid using Lipofectamine 3000. 24 h post transfection cells were either left untreated (DMSO control) or treated with 20 µM MG132 for another 4h and subjected to live cell confocal analysis after staining the nucleus with Hochest 33342. Doxycycline treatment induces both mCherry and sh-RNA expression. Scale bars, 5 µm. (D) A colony formation assay was conducted as described in the “Materials and Methods” section using a similar experimental setup in doxycycline inducible sh-Beclin1 stably expressing HEK293 cells transiently transfected with GFP-EBNA3C construct.

To further validate EBNA3C degradation induced upon MG132 treatment and its possible dynamics with autophagy machinery, live cell confocal microscopy analysis was conducted in doxycycline inducible sh-Beclin 1 stably expressing HEK293 cells transiently transfected with GFP-EBNA3C expressing construct (Fig. 4C). The results demonstrated a drastic reduction of overall EBNA3C puncta upon MG132 treatment as compared to DMSO control in the absence of doxycycline (active autophagy machinery) (Fig. 4C, upper panels). On the contrary, in the presence of doxycycline (Beclin 1 knockdown) MG132 treatment could not induce EBNA3C degradation as indicated by accumulation of GFP-EBNA3C puncta throughout the cell, predominantly in the cytoplasmic compartment (Fig. 4C, lower panels).

In order to assess whether or not proteasomal inhibition promotes EBNA3C degradation and thereby affecting its oncogenic potential, a colony formation assay was conducted using a similar experimental setup in doxycycline inducible sh-Beclin 1 stably expressing HEK293 cells transiently transfected with GFP-EBNA3C construct as described above (Fig. 4D). Indeed, the results demonstrated that impairment of autophagy pathway (Beclin 1 knockdown) partially reinstated EBNA3C colony formation ability when proteasome is blocked (Fig. 4D). Overall the results suggest that the increased degradation of EBNA3C complementing with reduced oncogenic potential observed with MG132 treatment was mediated at least in part by the autophagy-lysosomal pathway.

### EBNA3C interacts and colocalizes with p62-LC3B complex in the cytoplasmic fraction when proteasome is compromised

It has been demonstrated that p62/SQSTM1 directly binds to LC3B to facilitate degradation of ubiquitinated protein substrates by autophagy-lysosomal pathway [23]. To determine whether EBNA3C can participate within the p62-LC3B complex, HEK293 cells transiently transfected with either empty vector or flag-EBNA3C expressing construct, were subjected to immunoprecipitate with anti-flag monoclonal antibody (M2) with or without exposure of 20 µM MG132 for 4 h (Fig. 5A). The results demonstrated that EBNA3C forms a stable complex with both p62 and LC3B, irrespective of proteasomal activities (Fig. 5A). In order to validate the interaction of EBNA3C with p62-LC3B complex in a more endogenous setting, a similar immunoprecipitation experiment was carried out in LCLs (Fig. 5B). p62-LC3B complex was evidently precipitated with EBNA3C pull-down using a mouse monoclonal antibody (A10) in DMSO treated LCLs (Fig. 5B). However, owing to severe degradation of EBNA3C as well as p62 (increase of autophagy flux) in MG132 (1 µM, 12 h) treated LCLs resulted in lack of effective protein concentrations and accordingly no complex formation was detected when pull-down with anti-EBNA3C antibody as similar to mouse IgG control (Fig. 5B).

**Figure 5:**
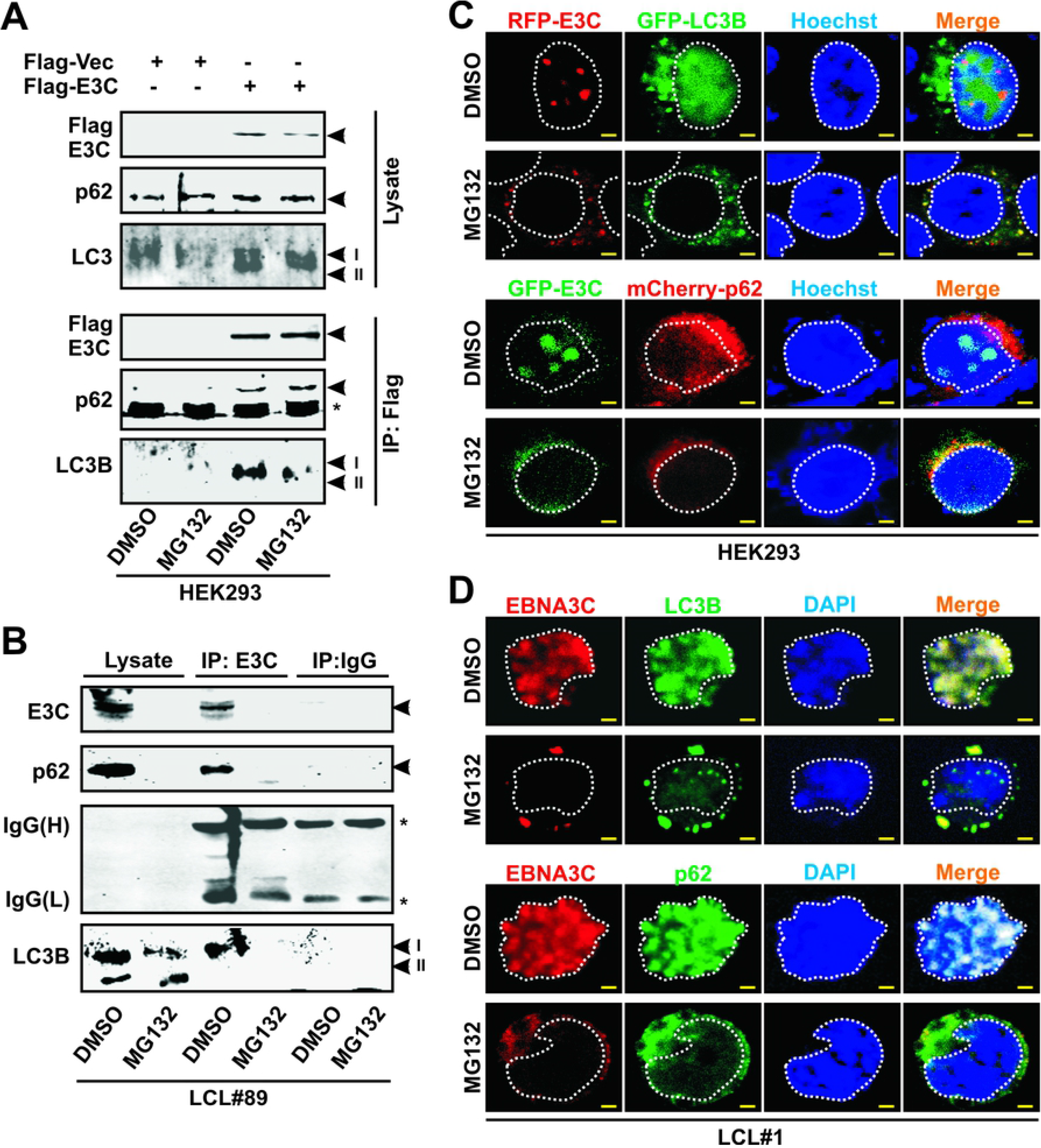
EBNA3C participates within the p62-LC3B complex when proteasome is inhibited. (A) ∼10 x 10^6^ HEK293 cells were transfected with either empty vector (pA3F) or flag-tagged EBNA3C expressing construct. 36h post-transfection cells were either left untreated (DMSO control) or treated with 20 µM MG132 for 4 h. After treatment, cells were harvested and subjected for immunoprecipitation with anti-flag antibody (M2). Precipitated products along with 10% whole cell lysates were run on gel and western blot analysis was performed with indicated antibodies. (B) ∼20 x 10^6^ LCLs (LCL#89) either left untreated (DMSO control) or treated with 1 µM MG132 for 12 h, were harvested and subjected for immunoprecipitation using EBNA3C specific mouse monoclonal antibody (A10). Rabbit anti-mouse IgG was used as isotype control. (C) ∼5 x 10^4^ HEK293 cells were co-transfected with the indicated expression plasmids using Lipofectamine 3000. 36 h post transfection, cells were further treated with DMSO or 20 µM MG132 for 4 h and subjected for live cell confocal analysis after staining the nucleus with Hochest 33342. (D) For confocal analysis in LCLs, ∼5 x 10^4^ LCLs (LCL#1) were treated with DMSO control or 1 µM MG132 for 12 h and subjected for immune staining with EBNA3C, p62, LC3B specific primary antibodies followed by incubation with Alexa Fluor conjugated secondary antibodies. Nuclei were counterstained using DAPI (4’,6’-diamidino-2-phenylindole) before mounting the cells. Each panel in (C-D) corresponds to single experiment of three independent experiments. Scale bars, 5 µm.

To corroborate the association of EBNA3C with p62-LC3B complex and degradation pattern in the absence or presence of MG132, colocalization experiments were carried out using both ectopic (HEK293) and endogenous (LCLs) expressing systems (Figs. 5C and 5D, respectively). First, HEK293 cells grown on glass bottom dishes were transiently transfected with RFP-tagged EBNA3C and GFP-tagged LC3B or GFP-tagged EBNA3C and mCherry-tagged p62 (Fig. 5C). Cells were further treated with 10 µM MG132 for 4h or left untreated (DMSO control) and subjected for live cell confocal microscopy analyses (Fig. 5C). The results demonstrated that MG132 treatment caused a drastic reduction of EBNA3C puncta formation in the nucleus compartments and increase in colocalization with both LC3B and p62 independently in the cytoplasmic fraction (Fig. 5C). A similar colocalization pattern of EBNA3C with LC3B and p62 was also observed in LCLs upon MG132 treatment (Fig. 5D). Overall, the results indicated that p62-LC3B binds to EBNA3C in physiological conditions, mediating its degradation via the autophagy-lysosomal pathway when proteasome is inhibited.

### EBNA3C is predominantly K63-linked polyubiquitinated when proteasome is inhibited

Studies suggested that proteins modified with K48-linked ubiquitin chains are targeted for the proteasomal pathway, whereas K63-linked chains can target substrates for degradation via autophagy [51, 52]. Previously it has been shown that EBNA3C can be polyubiquitinated at the N-terminal domain [28]. However, whether this polyubiquitination is K48-linked or K63-linked is still unclear. In addition, the UBA (ubiquitin associated) domain of p62 can bind to both K48- and K63-linked (with a higher affinity) polyubiquitin chains and regulate autophagic protein turnover [53]. These led us to further investigate the nature of polyubiquitinated chains associated with EBNA3C in the absence or presence of MG132. Immunofluorescence assays using specific antibodies against EBNA3C, total ubiquitin, K48- and K63-linked polyubiquitin chains in LCLs either left untreated (DMSO control) or treated with MG132, demonstrated that EBNA3C largely colocalized with ubiquitin irrespective of its type of chains within the nuclear compartments under physiological conditions (DMSO control), while MG132 treatment resulted in alteration of colocalization pattern from nuclear to predominantly cyctoplasmic along with drastic reduction of overall EBNA3C staining intensities (Fig. 6A). Using a similar experimental set up, western blot analyses of LCLs confirmed that MG132 treatment significantly enhanced both total ubiquitination and K48-linked polyubiquitination levels designated for proteasomal mediated degradation (Fig. 6B). In contrast, together with EBNA3C, K63-linked polyubiquitination level was notably decreased in MG132 treated LCLs, indicating that K63-linked polyubiquitinated substrates (including EBNA3C) are recycled in autophagy-lysosomal pathway, when proteasome is inhibited (Fig. 6B). Full blot analyses of EBNA3C (∼120 kDa) western blots using two different antibodies (mouse monoclonal followed by a sheep polyclonal) demonstrated the degradation/proteolytic cleavage pattern, identifying two predominant bands – one at ∼75 kDa and another at ∼45 kDa (Fig. 6B).

**Figure 6:**
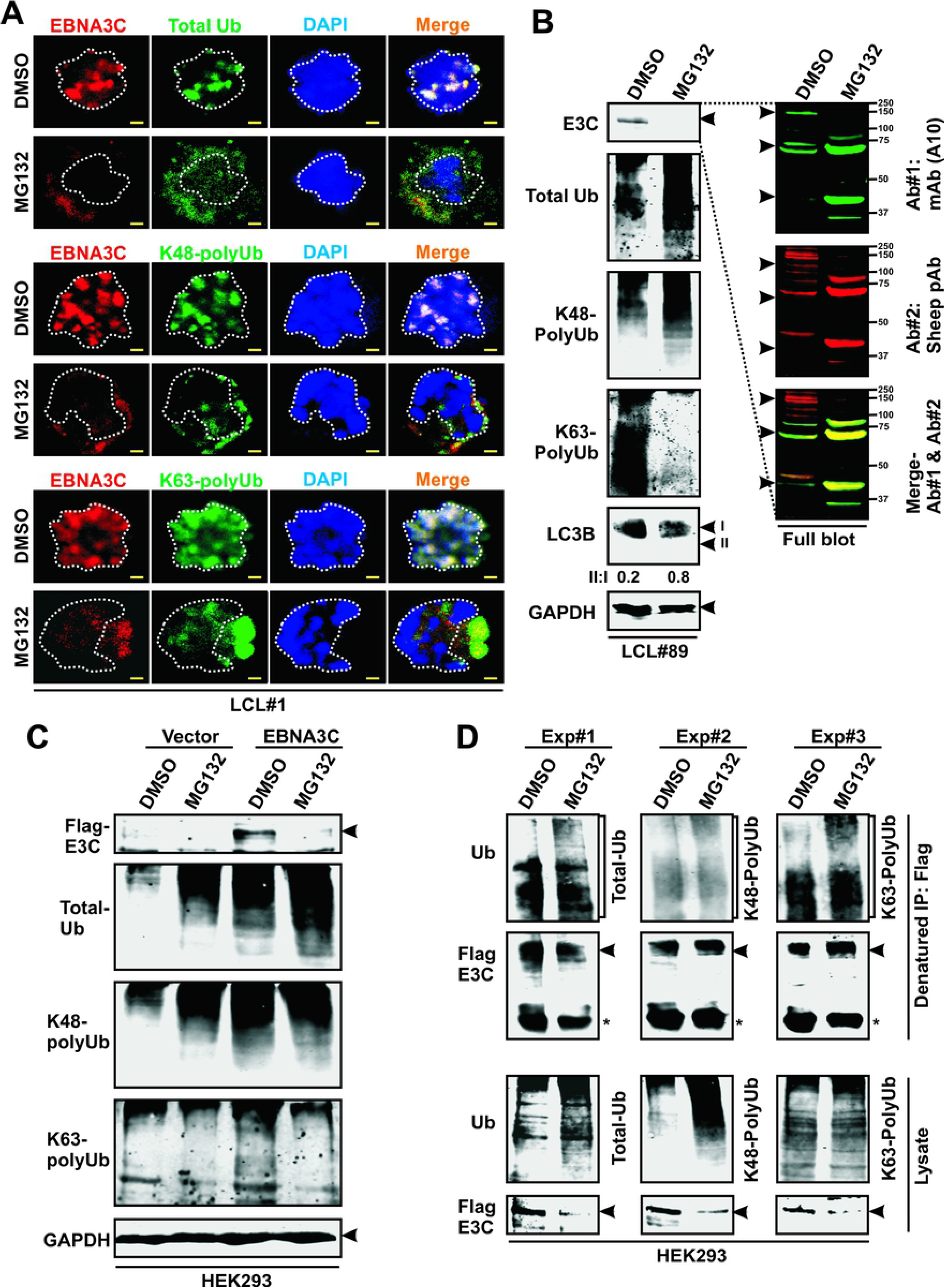
EBNA3C is predominantly K63-linked polyubiquitinated when proteasome is inhibited: LCLs were either left untreated (DMSO control) or treated with 1 µM MG132 for 12 h and subjected for (A) immunostaining with EBNA3C, total ubiquitin (Ub), K48- and K63-linked polyubiqitination specific antibodies; (B) western blot analyses with indicated antibodies. (A) Nuclei were counterstained with DAPI before mounting the cells for confocal analyses. (C-D) HEK293 cells transiently transfected with flag-tagged EBNA3C expression vector, either left untreated (DMSO control) or treated with 20 µM MG132 for 4h were subjected for (C) western blot analyses or (D) immunoprecipitation (IP) under denatured conditions using anti-flag antibody followed by western blot analyses using indicated antibodies. Each panel in (A) corresponds to single experiment of two independent experiments. Scale bars, 5 µm. Western blots in (B-C) were performed by stripping and reprobing the membranes with different antibodies.

To precisely determine EBNA3C specific poyubiquitnation pattern as well as to rule out the possible involvement of other viral oncoproteins, ectopic expression system was utilized where HEK293 cells were transiently transfected with either vector control or myc-tagged EBNA3C (Fig. 6C). The results demonstrated that presence of EBNA3C enhanced overall ubiquitination levels along with K48- and K63-linked polyubiqutinated chains in physiological conditions (DMSO control) as compared to empty vector (Fig. 6C). MG132 treatment further enhanced both total and K48-linked polyubiquitination, while significantly declined K63-linked polyubiquitination level (Fig. 6C), corroborating the previous experiments in LCLs. To validate EBNA3C polyubiquitination pattern, *in vivo* ubiquitination experiments was conducted in the absence and presence of MG132 in denaturing conditions allowing only covalent modifications (Fig. 6D). The results demonstrated that MG132 treatment specifically enhanced K63- but not K48-linked polyubiquitination of EBNA3C (Fig. 6D).

### N-terminal domain of EBNA3C is important for autophagy mediated degradation in response to proteasomal inhibition

Earlier demonstration of polyubiquitination at the N-terminal domain (residues 1-159) of EBNA3C [28] led us to further explore the domain(s) responsible for autophagy mediated degradation in response to proteasomal inhibition. To this end, HEK293 cells were transiently transfected with flag-tagged EBNA3C constructs expressing EBNA3C residues 1-365 (N-terminal), residues 366-620 (middle part) and residues 621-992 (C-terminal). As described earlier, 36h post-transfected cells were either left untreated (DMSO control) or treated with MG132 (20 µM) or CQ (50 µM) for another 4 h (Fig. 7A). The results demonstrated that only MG132 treatment caused significant reduction in expression of EBNA3C N-terminal domain as compared to DMSO or CQ treated cells (Fig. 7A). However, there was little or no change of ectopic expression levels of other two domains ranging from residue 366 to residue 992 in the absence or presence of both proteasomal and autophagy inhibitors (Fig. 7A). LC3II conversion confirmed autophagy activation, while enhanced polyubiquitination validated effective proteasomal inhibition upon MG132 treatment (Fig. 7A). *In vivo* ubiquitination experiments also validated polyubiquitination of EBNA3C at the N-terminal domain (residues 1-365) both in physiological conditions (DMSO control) and when proteasome is inhibited by MG132 (Fig. 7B).

**Figure 7:**
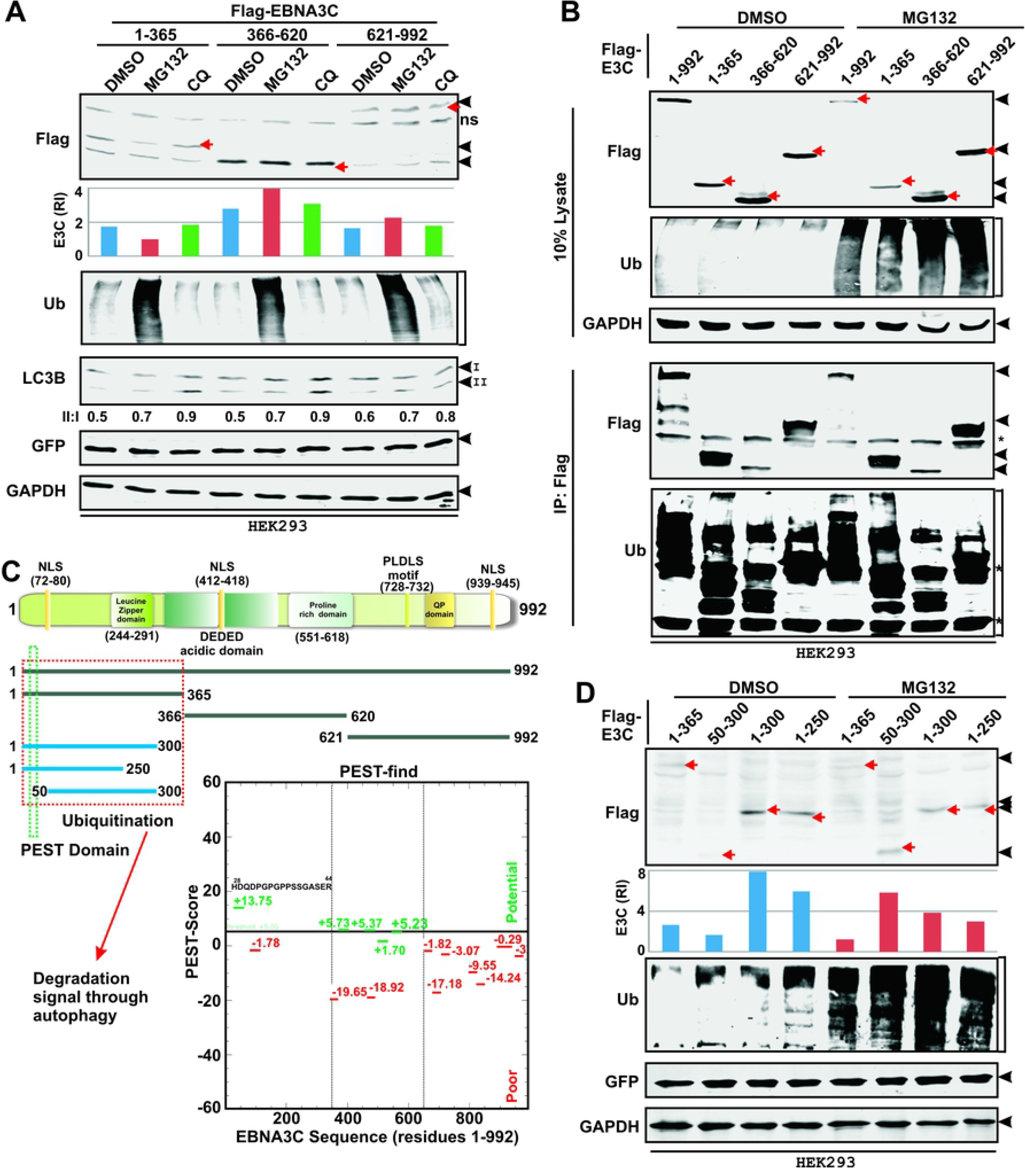
N-terminal domain of EBNA3C is important for autophagy mediated degradation in response to proteasomal inhibition. (A) HEK293 cells transiently transfected with plasmids expressing flag-tagged EBNA3C truncations – residues 1-365, 366-620 and 621-992, either left untreated (DMSO control) or treated with 20 µM MG132 or 50 µM Chloroquine (CQ), were harvested and subjected for western blot analyses with the indicated antibodies. (B) Using a similar experimental set up as described in (A), HEK293 cells transiently transfected with full-length (residues 1-992) and different domains of flag-tagged EBNA3C expression constructs, were subjected to immunoprecipitation (IP) with anti-flag antibody followed by western blot analyses with the indicated antibodies after stripping and reprobing the same membrane. (C) The schematic illustrates known structural motifs and different domains of EBNA3C that are used to determine residues important for autophagy mediated degradation upon proteasomal inhibition. Predicted PEST motifs were derived by using http://emboss.bioinformatics.nl/cgi-bin/emboss/epestfind (green: potential; red: poor). (D) HEK293 cells transiently transfected with expression plasmids for flag-tagged EBNA3C N-terminal trunactions (residues 1-365, 50-300, 1-300 and 1-250) either were subjected to western blot analyses with the indicated antibodies after a similar treatment as described in (B). Protein band intensities in (A and D) were quantified by Odyssey imager software and indicated either as bar diagrams or LC3-II/I ratio at the bottom of each corresponding lane. Representative gel pictures are shown of two independent experiments. GAPDH and GFP blots were performed as loading and transfection efficiency controls, respectively.

To identify whether the N-terminal domain contains any proteolytic cleavage site, full length EBNA3C sequence was searched for the presence of putative PEST (proline, glutamic acid, serine and threonine) sequences. Near the N-terminal domain (residues 28-44) a potential PEST sequence ‘HDQDPGPGPPSSGASER’ was identified (Fig. 7C). In order to determine whether this identified PEST sequence is responsible for autophagy dependent degradation in response to proteasomal inhibition, three more flag-tagged EBNA3C truncated constructs were generated expressing residues 50-300 (lacking PEST motif), residues 1-300 and residues 1-250. HEK293 cells transiently transfected with these constructs along with EBNA3C residues 1-365 expressing plasmid were subjected to western blot analyses after treatment with either DMSO control or MG132 (Fig. 7D). While MG132 treatment caused obvious stabilization of residues 50-300, significant reduction of protein expression was observed when first 50 residues were included in all three EBNA3C truncations (Fig. 7D). Taken together, the results indicated that EBNA3C N-terminal first 50 residues containing PEST sequence might be responsible for autophagy-lysososmal mediated degradation when proteasome is inhibited.

### EBNA3C N-terminal domain is degraded in the both nuclear and cytoplsamic fractions upon proteasomal inhibition

In order to continue exploring the mechanisms involved in EBNA3C degradation, full length EBNA3C sequence was investigated for potential leucine-rich nuclear export signal (NES) motifs in NetNES 1.1 Server [54]. Two predicted NES sequences of EBNA3C were determined near the N-terminal (residues 124-132 “LQALSNLIL”) and C-terminal (residues 862-869 “LQLSLVPL”) regions (Fig. 8A). However, interestingly, no predicted NES sequences were found for both EBNA3A and EBNA3B proteins. To confirm this prediction, a subcellular fractionation was performed using HEK293 cells transiently transfected with myc-EBNA3C (Fig. 8B). The results demonstrated that EBNA3C was indeed fractionated in both nuclear and cytoplasmic sections and upon MG132 treatment EBNA3C was specifically found to be degraded in the cytoplasmic fraction (Fig. 8B). However, in contrast, p53 degradation occurred in the nuclear fraction in response to MG132 treatment (Fig. 8B). The efficiency of subcellular fractionation was determined by GAPDH and lamin A/C blots as cytoplasmic and nuclear reference proteins, respectively (Fig. 8B). To further correlate subcellular localization pattern and proteasomal inhibition mediated EBNA3C degradation, live-cell confocal microscopy analysis was carried out in HEK293 cells transiently transfected with GFP-tagged EBNA3C (Fig. 8C). MG132 treatment resulted in co-localization of EBNA3C (wild-type, residues 1-992) with the lsyosomal fraction in the cytoplasm (stained with LysoTracker Red DND-99) (Fig. 8C).

**Figure 8:**
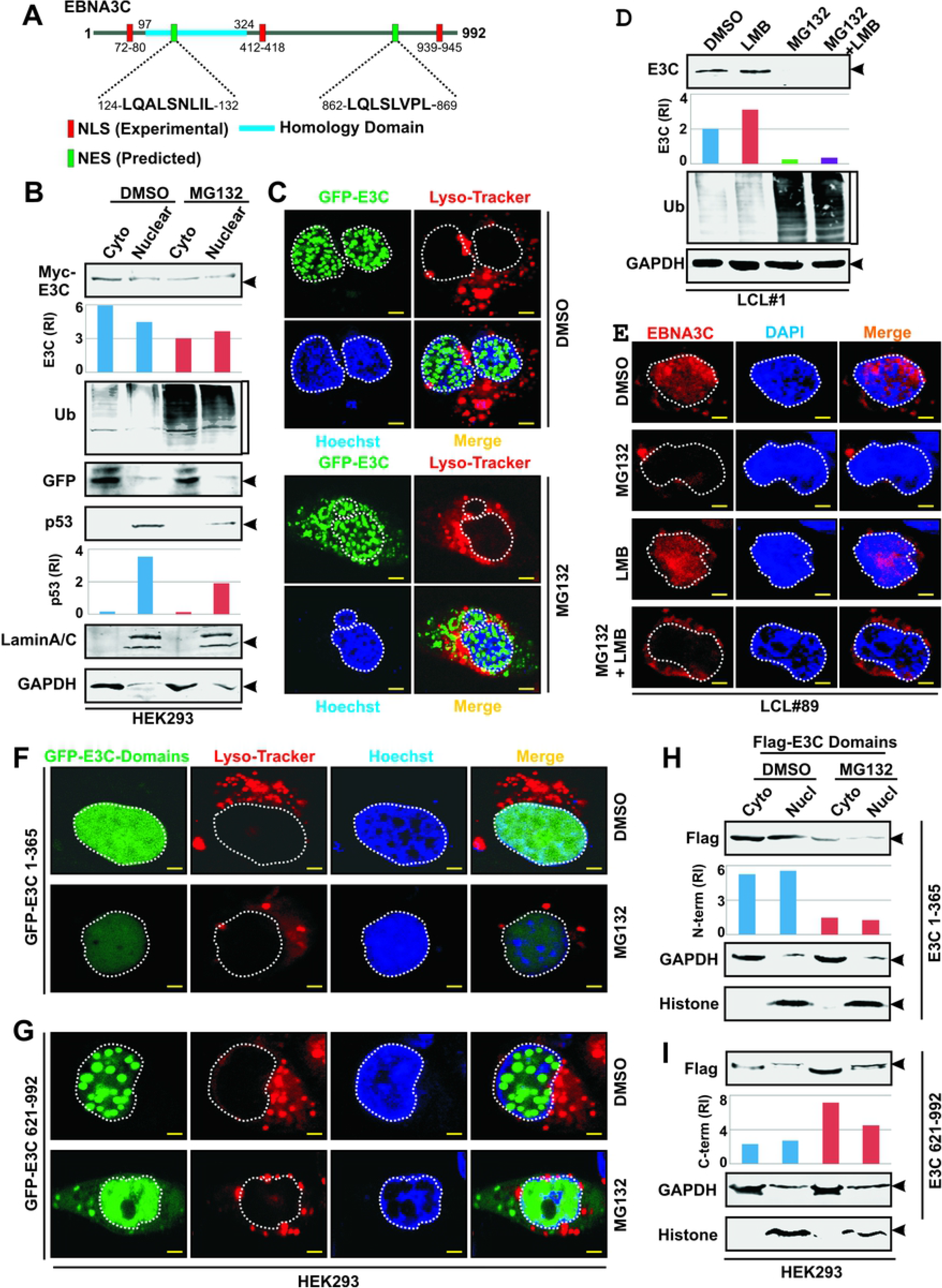
EBNA3C N-terminal domain is degraded in both nuclear and cytoplsamic fractions upon proteasomal inhibition. (A) Schematic shows the predicted nuclear export signal (NES) sequences of EBNA3C, derived from http://www.cbs.dtu.dk/services/NetNES/. (B-C) HEK293 cells were transiently transfected with the indicated expression plasmids. 36 h post-transfection cells were further treated with DMSO control or 20 µM MG132 for 4 h and subjected to (B) subcellular fractionation followed by western blot analyses using the indicated antibodies and (C) live cell confocal analyses. (D-E) LCLs were either left untreated (DMSO control) or treated with 0.5 µM MG132, 20 ng/ml leptomycin B (LMB) or MG132 plus LMB. 24 h post-treatment cells were subjected for either (D) western blot analysis or (E) immunostaining with the indicated antibodies. (C, F-G) Nuclei and lysosomes were counterstained with Hoechst 33342 and LysoTracker Red DND-99, respectively. Nuclei in (D) were counterstained by DAPI before mounting the cells. Each panel of confocal images in (C, E-G) is representative pictures of two independent experiments. Scale bars, 5 µm. (H-I) HEK293 cells transiently transfected with plasmids expressing flag-tagged EBNA3C N-terminal (residues 1-365) and C-terminal (residues 621-992) domains in a similar experimental set up as described in (B) were subjected to subcellular fractionation as per Manufacturer’s instruction, followed by western blot analyses with the indicated antibodies. (B and H-I) GAPDH was used as reference protein for cytoplasmic fraction, while lamin A/C (B) and histone (H-I) blots were performed as nuclear reference proteins. Protein bands in (B, D, H-I) were quantified by Odyssey imager software and indicated as bar diagrams at the bottom of corresponding lanes.

Leptomycin B, an inhibitor of nuclear export, can cause the nuclear accumulation of proteins that shuttle between the cytosol and nucleus by targeting CRM1 (exportin 1), an evolutionarily conserved receptor for the nuclear export signal of proteins [55]. To further establish the role of nuclear export signal in mediating EBNA3C degradation in the cytoplasmic compartment, LCLs were treated with MG132 in the presence and absence of leptomycin B (20 ng/ml) and subjected for western blot and confocal analyses (Figs. 8D and E, respectively). While leptomycin B treatment alone slightly enhanced EBNA3C expression in LCLs, it could not manage EBNA3C degradation induced by proteasomal inhibition (Figs. 8D-E). The result suggests that EBNA3C degradation might occur in both nuclear and cytoplasmic compartments or leptomycin B failed to block cytoplasmic export of EBNA3C completely. The effect of leptomycin B was tested by conducting a fractionation experiment using HEK293 cells transiently transfected with flag-tagged EBNA3C with or without 24 h leptomycin B post-treatment (Fig. S2). The results demonstrated that leptomycin B treatment significantly enhanced (∼3 fold) EBNA3C amount in nuclear fraction when compared to DMSO control (Fig. S2). The efficiency of fractionation was determined by western blot analyses against GAPDH as cytoplasmic protein and histone as nuclear protein (Fig. S2).

In order to determine whether both predicted NES sequences are functional or one of them are actively engaged in EBNA3C degradation upon proteasomal inhibition, GFP-tagged EBNA3C N-terminal (residues 1-365) and C-terminal (residues 621-992) domains containing two predicted NES sequences were utilized (Figs. 8F-I). Live cell confocal experiments demonstrated that while the overall fluorescence intensity of N-terminal domain was drastically reduced particularly in the nuclear compartment, there was a significant increase in cytoplasmic translocation of C-terminal domain (compare Figs. 8F and G). Moreover, the results demonstrated that the puncta pattern of EBNA3C conferred by the C-terminal domain, while the N-terminal domain presented rather a diffused pattern of fluorescence (Figs. 8F-G). To get a clearer picture, subcellular fractionation was carried out in HEK293 cells transiently transfected with both flag-tagged N-terminal and C-terminal domains of EBNA3C with or without MG132 treatment (Figs. 8H and I, respectively). As similar to the full-length EBNA3C, both N-terminal and C-terminal domains were almost equally distributed in nuclear and cytoplasmic fractions (Fig. 8H-I). Moreover, as previously observed, p53 was only degraded in the nuclear fraction in these experiments (data not shown). The efficiency of subcellular fractionation was similarly validated by GAPDH and histone blots as cytoplasmic and nuclear reference proteins, respectively (Fig. 8H-I). Quantification of the band intensities clearly indicated that MG132 treatment led to degradation of the N-terminal domain in both nuclear and cytoplasmic fractions, whereas the C-terminal domain was somewhat stabilized in both fractions (Fig. 8H-I).

Taken together, the results suggest that nuclear export signal residing at both N-terminal and C-terminal domains play a crucial role for EBNA3C degradation in the cytoplasm through autophagy mechanism when proteasome is inhibited or overload ded due to accumulation of mis-/un-folded proteins. Because the autophagy mediated degradative signal is located at the N-terminal region of EBNA3C (residues 1-50), to determine whether the N-terminal domain is also responsible for forming complex with autophagy components - p62/LC3B, an immunoprecipitation experiment was performed in HEK293 cells transiently transfected with either vector control or flag-tagged EBNA3C truncations – N-terminal (residues 1-365), middle part (residues 366-620) and C-terminal (residues 621-992) (Fig. S3A). The results demonstrated that the only N-terminal domain bound to both p62 and LC3B (Fig. S3A-D), which likely initiates EBNA3C degradation via autophagy-lysosomal pathway.

### Caspases and reactive oxygen species do not affect proteasome inhibitor mediated degradation of EBNA3C

Studies suggest that caspases together with autophagy and UPS also contribute in proteolytic mechanisms particularly during apoptosis [19, 56]. In addition, proteasome inhibitors have been shown to induce apoptotic cell death through the formation of reactive oxygen species (ROS) [56]. Apoptotic induction due to MG132 treatment, further prompted us to investigate whether caspases or ROS are involved in MG132 induced EBNA3C’s degradation. However, no potential cut sites on EBNA3C’s sequence were found for Caspases 1-10 using a web based peptide cutter tool (https://web.expasy.org/peptide_cutter/). Western blot results and confocal analyses also demonstrated that addition of a pan-caspase inhibitor, Z-VAD(OMe)-FMK could not rescue EBNA3C degradation in response to proteasomal inhibition (Fig. S4A-B).

To evaluate the role of ROS in EBNA3C degradation N-acetyl-L-cysteine (NAC), a known ROS scavenger [57], was used and tested its ability to antagonize MG132 mediated ROS generation in LCLs (Fig. S5A). However, blocking the ROS by NAC did not restore EBNA3C expression as shown by both western blot and confocal analyses (Fig. S5B-C). Overall, the data suggest that proteasomal inhibition mediated apoptotic induction and ROS had limited effect on modulation of EBNA3C expression. A similar result was also obtained for p53 degradation (Figs. S4A and S5A).

### Proteasome inhibitors can be used as potential therapeutic strategy against EBV associated B-cell lymphomas

Proteasome inhibitors specifically inhibit degradation of poly-ubiquitinated proteins through UPS, resulting in the stabilization of several tumor suppressor proteins that eventually promote cell-cycle arrest and apoptosis [58, 59]. Previously it has been demonstrated that bortezomib, the first FDA approved proteasome inhibitor for the treatment of myeloma and mantle cell lymphoma, promotes EBV lytic cycle activation in Burkitt’s lymphoma (BL) [50]. EBNA3C degradation induced by proteasomal inhibition prompted us to evaluate proteasome inhibitors as a potential therapeutic strategy against EBV associated B-cell lymphomas where EBNA3C is expressed, particularly those are generated in an immunocompromised background. Soft agar based colony formation assays and cell viability experiments using trypan blue exclusion method on two LCLs (LCL#1 and LCL#89) demonstrated that both laboratory based and clinically approved proteasome inhibitors, MG132 and bortezomib, respectively, caused a significant reduction of total number of viable cells and colony formation ability in a dose dependent manner (Fig. 9A-D). Bortezomib (LD_50_: ∼0.5 µM) showed approximately 10 fold higher cytotoxity than that of MG132 (LD_50_: ∼5 µM) in LCLs (Fig. 9D). To determine whether proteasome inhibitors may exert different cytotoxic effects on B-cell lymphomas in respect to EBV infection status, cell viability assays was conducted in two EBV negative (BJAB and DG75) and two EBV positive (Namalwa and Raji) BL lines (Fig. 9E). The results demonstrated no significant variation of proteasome inhibitors induced cell death between EBV negative and EBV positive lines (Fig. 9E). Since EBNA3C was shown to enhance cytoprotective autophagy as well as block apoptosis in response to several cellular insults, similar cell viability assays were carried out in both EBNA3C stably expressing BJAB cells and EBNA3C knockdown LCLs to specifically determine if EBNA3C expression may influence proteasome inhibitors mediated cell death (Fig. 9F and G, respectively). The results demonstrated that while EBNA3C knockdown LCLs were more susceptible to cell-death induced by proteasome inhibitors, EBNA3C expression in B-cells led to a significant protection as compared to the control lines (Fig. 9F-G). EBNA3C expression was checked in different cell lines used in this study by western blot analyses (Fig. S6). Overall, the data show promise for proteasome inhibitors as anti-cancer drugs against multiple EBV associated B-cell lymphomas irrespective of EBNA3C expression.

**Figure 9:**
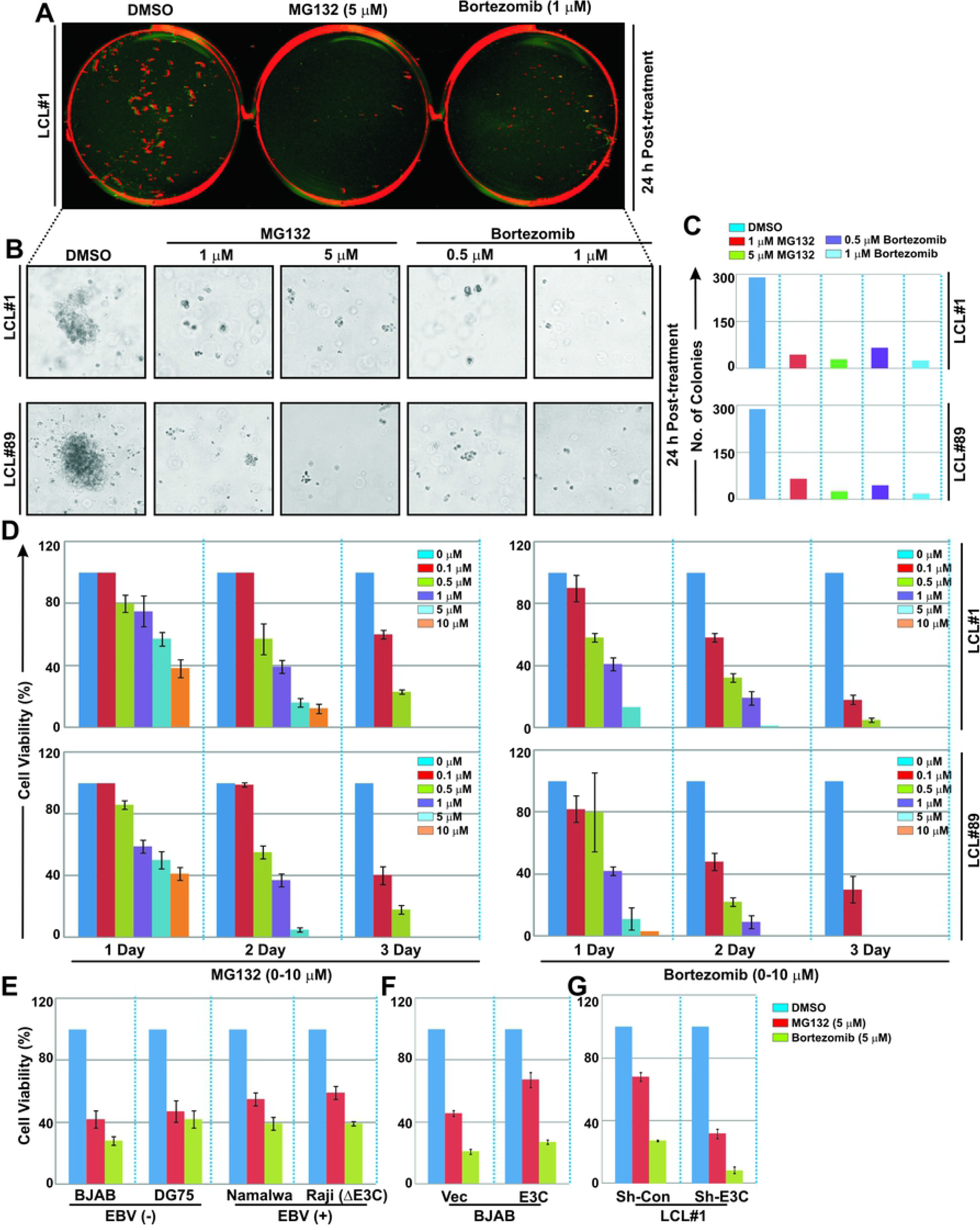
Proteasome inhibitors can be used as potential therapeutic strategy against EBV associated B-cell lymphomas. (A-C) 1 × 10^5^ LCLs (LCL#1 and LCL#89) were either left untreated (DMSO control) or treated with increasing concentrations of MG132 (1-5 µM) or bortezomib (0.5-1 µM) for 24 h and subjected for soft agar colony formation assay as described in the “Materials and Methods” section. After 14 days colonies were stained with 0.1% crystal violet and scanned using Odyssey CLx Imaging System and the number of colonies were measured by Image J software and plotted as bar diagrams in (C). Prior to staining, each well was also photographed (bright-field) using a Fluorescent Cell Imager as shown in (B). (D-G) ∼0.5 × 10^5^ cells plated into each well of six-well plates were either left untreated (DMSO control) or treated with increasing concentrations (0-10 µM) of MG132 or bortezomib for the indicated time points at 37^0^C in a humidified CO_2_ chamber. Viable cells from each well were measured by Trypan blue exclusion method using an automated cell counter. Error bars represent standard deviations of duplicate assays of two independent experiments.

## Discussion

EBNA3 family proteins consisting of EBNA3A, EBNA3B and EBNA3C are three closely related EBV nuclear proteins that are expressed only in latency III associated B-cell lymphomas and *in vitro* transformed continually proliferating LCLs [13, 14]. Genetic analysis using recombinant viruses has shown that EBNA3A and EBNA3C, but not EBNA3B, are essential for B-cell activation and subsequent immortalization *in vitro* [8,9,45]. Previously, these EBNA3 proteins were shown to be extremely stable in growing LCLs, although *in vitro* they interact with 20S proteasome, one of the major proteolytic components of 26S proteasome in ubiquitin-proteasome system (UPS) of eukaryotic cells [33]. 26S proteasome, a 2.5-MDa self-compartmentalized proteolytic machine located in the cytosol and nucleus, comprises of two 19S regulatory particles responsible for recognition of poly-ubiquitinated proteins marked for proteolytic degradation, assemble each end of the barrel-shaped 20S proteasome, where proteolysis takes place [60]. In eukaryotic cells, UPS is considered as one of the key protein degradation mechanisms, affecting an array of important cell pathways [20, 21]. Consequently, specific chemical inhibitors of UPS have recently emerged as attractive anticancer drugs [58, 59]. For example, bortezomib (Velcade) has been approved by the US Food and Drug Administration (FDA) as the first proteasome inhibitor for the treatment of refractory multiple myeloma and mantle cell lymphoma [46, 61].

Proteasome function can be hindered in many ways. For example, cancer cells for its high metabolic activity often encounter ER stress due to accumulation of misfolded and aggregated proteins exerting an unfolded protein response (UPR) [62]. Such protein aggregates can bind and stabilize an inactive closed conformation of the 26S proteasome [63]. Additionally, UPS alone is not capable enough to alleviate UPR and resulting in apoptotic cell death. Thus, in order to survive and properly function, cells must obliterate these protein aggregates, where autophagy-lysosomal pathway comes into the picture as a compensatory proteolytic mechanism particularly when proteasome function is inhibited [64, 65]. In agreement to this, a growing body of evidence demonstrates proteasomal inhibition can promote degradation of a number of important cellular proteins thorough activating autophagy mechanism. For example, it has been shown that while wild-type p53 is stabilized by autophagy activation, oncogenic mutant p53 is degraded by autophagy-lysosomal pathway in response to proteasomal inhibition in breast cancer cell lines [37]. Other examples of cell oncoproteins that are degraded through autophagy include the two fusion oncoproteins - BCR-ABL and PML-RARA, anterior gradient 2 (AGR2), a member of the protein disulfide isomerase (PDI) family, and cyclin dependent kinase 1 (CDK1) [38,66,67,68]. In addition, JC virus (JCV) encoded large T-antigen was shown to be degraded through autophagy mediated by Bag3, a member of the Bcl-2-associated athanogene (Bag) family of proteins [69]. Current knowledge about the role of autophagy in regulating expressions of EBV encoded oncoproteins is primarily limited to EBNA1 and LMP1. While LMP1 modulates autophagy pathway to regulate its own degradation, EBNA1 is degraded through autophagy-lysosomal pathway for MHC class II mediated antigen presentation [34, 70]. Thus, the available data are consistent with a notion that disruption of autophagy function can act as a tumor barrier through degradation of both cell and viral oncoproteins in response to various cellular insults.

In this report, for the first time we show that proteasomal inhibition enhances degradation of a viral oncoprotein, EBV encoded essential oncoprotein EBNA3C through autophagy-lysosomal pathway (Fig. 10). Although share a considerable structural similarities [16], proteasomal inhibition causes degradation of only two EBNA3 family members oncoproteins – EBNA3A and EBNA3C, but not the third member – the viral tumor suppressor protein EBNA3B. Recently we have shown that EBNA3C enhances the basal level of autophagy, which serves as a prerequisite for EBNA3C mediated cell proliferation and apoptotic inhibition [36]. In the absence of growth promoting signals this autophagy activation is further enhanced through engaging an epigenetic regulation [36]. It is not entirely clear to what extent basal autophagy contributes to EBNA3C degradation, while autophagic disruption arises when autophagy is stimulated higher than the basal levels in response to stress signals (including UPR or starvation) or drug treatment (such as proteasome inhibitors). As discussed above, autophagy can also be induced in response to a heavy burden of aggregated proteins [64, 65]; in our model autophagy induction alone was unable to induce EBNA3C degradation. Dual treatment with inhibitors of the UPS (MG312) and autophagy (chloroquine treatment or sh-RNA mediated Beclin1 knockdown) revealed that autophagy-lysosomal pathway plays a crucial role in degradation of EBNA3C (and possibly along with other protein aggregates) caused by UPS inhibition. Our data also indicate that autophagy inhibition could not fully compensate EBNA3C expression caused by UPS impairment. Interestingly, it has earlier been shown that basal but not enhanced autophagy activity regulates UPS mediated Sirt3 degradation in chronic myeloid leukemia cells [71]. This offers an adaptive mechanism by which autophagy in association with UPS mitigates oxidative stress in the leukemia cells.

**Figure 10:**
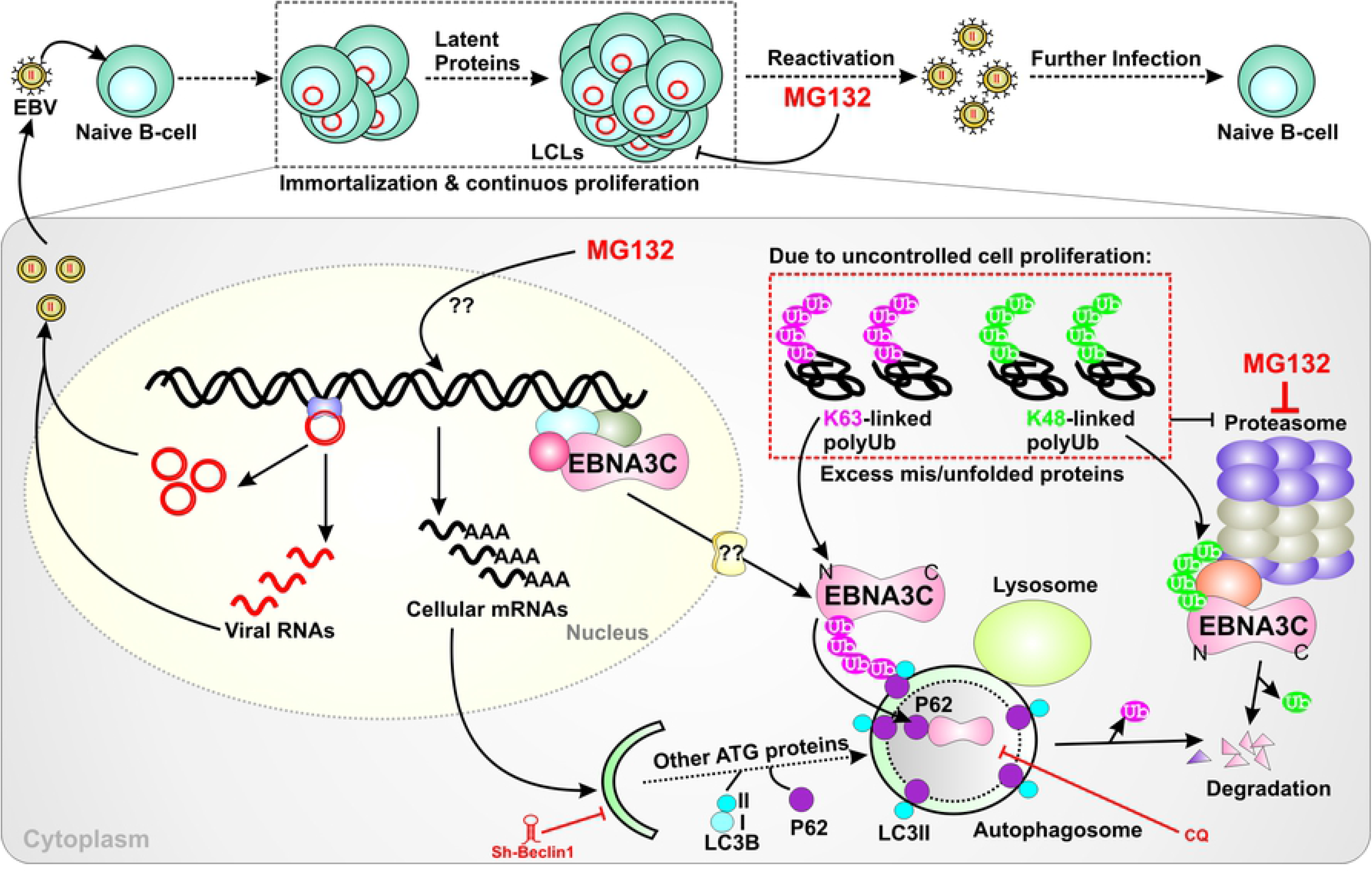
Schematic representation of proteasomal inhibition mediated EBNA3C’s degradation. Due to uncontrolled cell proliferation, cancer cells often encounter excess mis-/un-folded protein aggregates, which are subsequently labeled with either K48-linked or K63-linked polyubiquitination for proteolytic degradation. While K48-linked ubiquitin chains are targeted for the proteasomal pathway, K63-linked directs autophagy mediated protein degradation. Upon proteasomal inhibition, EBV oncoprotein EBNA3C is predominantly tagged with K63-linked polyubiquitin chains, translocated to cytoplasm by an unknown mechanism and degraded through autophagy-lysosomal pathway. The N-terminal domain plays a central role in EBNA3C’s degradation through autophagy mechanism by participating within the p62-LC3B complex. Suppression of autophagy pathway (Beclin1 knockdown or chloroquine, CQ treatment) reverses EBNA3C degradation in response to proteasomal inhibition. Additionally, proteasomal inhibitors (such as MG132) induce both autophagy and viral gene transcription that eventually activate viral lytic replication. The results provide foundation to exploit proteasome inhibitors as potential therapeutic approach for EBV associated B-cell lymphomas, where EBNA3C is expressed, typically diagnosed in immunocompromised individuals.

Earlier, we have demonstrated that EBNA3C expression increases basal level of K63-linked polyubiquitination, which further increases in absence of growth promoting signals [36]. Additionally, EBNA3C was shown to recruit UPS to regulate turn-over of many important cell oncoproteins and tumor suppressors [27,28,29,30,31,72,73,74]. Initially Knight et al. demonstrated the poly-ubiquitination on the N-terminal domain of EBNA3C [28]. However, the authors conducted the study in native conditions and thus cannot rule out the possibility of association of other ubiquitinated proteins within the complex. In this study, using immunoprecipitation under denaturing conditions we now show for the first time that EBNA3C are specifically covalently tagged by K63-linked polyubiquitin chains. Ubiquitin chains with these different linkages are thought to direct proteins towards different fates owing to the specificity of binding partners for specific lysine-linkages. While K48-linked polyubiquitin chains are coupled with proteasome mediated degradation, K63-linked chains direct degradation of substrate protein through autophagy-lysosomal pathway [51, 52]. The presence of K63-chain on EBNA3C offers a model in which K63–ubiquitin labeled EBNA3C is targeted for proteolytic degradation through autophagy-lysosomal pathway. The autophagic adaptor protein p62/SQSTM1 selectively binds to K63-linked polyubiquinated chains and subsequently recruits to LC3-positive autophagosomes for degradation [23, 52]. Our data also reveals that EBNA3C forms a stable complex with p62/LC3B. Bioinformatics analyses could not however reveal any putative LIR (LC3 interacting region) domain in EBNA3C [24, 75], suggesting that EBAN3C perhaps does not directly bind to LC3B. The interaction may occur through p62 and K63-polyubiquition linked EBNA3C.

The N-terminal domain of EBNA3C is of particularly interesting, at least in the context of interacting with plethora of important cell proteins particularly involved in cell-cycle activation and blockade of apoptosis [14,30,31,72,74,76,77,78]. Interestingly, many of these proteins’ turn over or stability is deregulated through UPS specifically recruited by the N-terminal domain of EBNA3C. For example, EBNA3C enhances degradation of p27^KIP1^ and p53 tumor suppressor proteins through recruiting SCF^Skp2^ and Mdm2 E3 ligase activities, respectively and thereby promoting EBV induced B-cell lymphomagenesis [28, 30]. Moreover, using genetic engineering the N-terminal residues were shown to be essential for survival of EBV transformed B-lymphocytes [79]. Our results demonstrate that this N-terminal domain (residues 1-365) participates within the p62-LC3B complex and particularly the first 50 residues of EBNA3C comprising a putative PEST domain is responsible for autophagy mediated degradation upon proteasomal inhibition. This supports a model whereby the N-terminal domain plays a feedback mechanism to control EBNA3C expression through autophagy-lysosomal pathway in response to proteasomal impairment. The presence of putative nuclear export signal (NES) sequences in both N- and C-terminal domains also indicate a possible mechanism of EBNA3C degradation through autophagy in the cytoplasmic fraction. In fact, our results demonstrate that MG132 treatment leads to EBNA3C’s degradation in both nucleus and cytoplasm, whereas p53 specifically degrades in the nuclear compartments. However, the precise mechanism by which EBNA3C shuttles between nucleus and cytoplasm in response to proteasomal inhibition is yet to be defined.

Previously, proteasome inhibitors, such as MG132 have been demonstrated to exert substantial protective effect on cardiovascular and renal injury and appear to be potentially effective drugs in blocking oxidative damage as well as increased atherosclerosis in rabbits [80,81,82,83]. MG132 treatment was shown to enhance progerin clearance through autophagy and regulating splicing mechanisms by downregulating SRSF-1 [39]. Besides its proteolytic functions, proteasomal machinery also provides non-proteolytic activities engaging transcriptional and post-transcriptional regulation of multiple cellular genes [39,66,84]. In agreement to this, our RNA-seq data reveals that proteasomal inhibition enhances transcriptional activation of genes (viz. MAP1LC3B, p62/SQSTM1 and GABARAPL1) involved in autophagosome formation. Moreover, in contrast to proteolytic degradation of EBNA3 proteins, MG132 treatment results in transcriptional activation of both EBV latent including the EBNA3 family genes and lytic genes, which subsequently promotes viral lytic cycle. There is a growing interest in pharmacologic activation of lytic viral gene expression as a potential therapeutic strategy against several virus associated human cancers [48]. Of note, it has been earlier demonstrated that bortezomib treatment augments EBV lytic cycle activation in Burkitt’s lymphoma cell lines [50]. The authors suggested that UPR regulator CCAAT/enhancer-binding protein (CEBPβ), a member of the bZIP family of transcription factors, transcriptionally activates BZLF1 or ZTA expression that eventually controls EBV lytic cycle activation [50]. In this line, our lab is currently investigating the role of UPR activators in EBNA3C degradation as a prospective therapeutic option. Altogether these results suggest that proteasome inhibitors can have encouraging therapeutic effects without having any potential cytotoxicity.

In conclusion, herein the results provided the basis for the development of novel class of drug against multiple EBV linked B-cell lymphomas, expressing EBNA3 family proteins. We provided evidence that proteasomal inhibition specifically promotes degradation of two viral oncoprotein EBAN3A and EBAN3C through autophagy-lysosomal pathway along with induction of viral lytic cycle. A new promising path hence begins for therapeutic development of proteasome inhibitors (such as bortezomib) in combination with EBV replication inhibitors (such as Ganciclovir) toward a better treatment in near future for patients affected with EBV associated B-cell lymphomas particularly those generated in an immunocompromised background.

## Materials and Methods

### Drugs

MG132 (ab141003), bortezomib (ab142123) and Z-VAD(OMe)-FMK (ab120487) were obtained from Abcam (Cambridge, UK). Chloroquine diphosphate salt (C6628), Cyclohexamide (C7698), Leptomycin B (L2913), N-Acetyl-L-cysteine (A9165), Doxycycline hyclate (D9891), G418 sulfate (345810), 12-O-Tetradecanoylphorbol-13-acetate or TPA (P1585), sodium butyrate (B5887), Polybrene (TR-1003-G) and Puromycin dihydrochloride (P8833) were brought from Sigma-Aldrich Corp. (St. Louis, MO, USA).

### Plasmids

Myc-, flag-, GFP- and RFP-tagged EBNA3C constructs in pA3M, pA3F, pEGFP-C1 and pDsRED-Monomer-N1 vectors, respectively were previously described [27,29,36]. Myc-tagged EBNA1 expressing construct in pA3M was described earlier [85]. Myc- and flag-tagged EBNA3C truncation constructs expressing EBNA3C residues 1-365, 366-620 and 621-992 were previously described [27, 29]. Myc-tagged EBNA3C N-terminal residues 1-365 expressing construct was used as a template for preparing three more EBNA3C truncations – residues 1-300, 1-250 and 50-300 in pA3M (pCDNA3.1-3X Myc) at *EcoR*I and *Not*I restriction sites. pA3M-EBNA3C construct was used as a template to clone GFP-tagged EBNA3C truncation constructs expressing residues 1-365 and 621-992 in pEGFP-C1 vector at *EcoR*I and *Sal*I restriction sites. All the constructs were further verified by Sanger dideoxy based DNA sequencing (Eurofins Genomics India Pvt. Ltd., India). pEGFP-C1 and pDsRED-Monomer-N1 plasmids were obtained from Clontech Laboratories, Inc. mCherry-tagged p62 expressing plasmid was a kind gift from Edward M. Campbell (Loyola University Chicago, IL, USA). pBabePuro-GFP-LC3 construct was a kind gift from Jayanta Debnath (Addgene plasmid# 22405). pTripz-mCherry-Sh-Beclin1 construct (Clone ID: V2THS_23692) was obtained from GE Healthcare Dharmacon Inc. (Lafayette, CO, USA). pGipz-Sh-Control and pGipz-Sh-EBNA3C were previously described. Lentiviral packaging vectors - pCMV-VSV-G (Addgene plasmid# 8454), pRSV-Rev (Addgene plasmid# 12253), and pMDLg/pRRE (Addgene plasmid# 12251) were kind gifts from Robert A. Weinberg (Whitehead Institute for Biomedical research, Cambridge, MA, USA).

### Antibodies

Mouse monoclonal antibody against EBNA3C (A10) generated from hybridoma cells was previously described. Sheep polyclonal antibodies against EBNA3A (ab16126), EBNA3B (ab16127) and EBNA3C (ab16128) were bought from Abcam (Cambridge, UK). Rabbit polyclonal anti-GFP (ab290), rabbit anti-Sheep IgG (H+L), rabbit polyclonal anti-Histone H3 (ab1791) and rabbit monoclonal anti-LMP1 (ab136633) antibodies were also obtained from Abcam (Cambridge, UK). Mouse monoclonal antibodies directed against p62/SQSTM1 (Clone# 3/P62 lck ligand), Beclin1 (Clone# 20/Beclin), cleaved PARP (Clone# Asp214), total ubiquitin (Clone# 6C1.17) were purchased from BD Biosciences (Franklin Lakes, NJ, USA). Rabbit monoclonal antibodies directed against LC3A/B (Clone# D3U4C), K48-linkage specific polyubiquitin (D9D5) and K63-linkage specific polyubiquitin (D7A11) were purchased from Cell Signaling Technology Inc. (Danvers, MA, USA). Mouse monoclonal anti-flag (M2) and anti-cMyc (9E10) antibodies were purchased from Sigma-Aldrich Corp. (St. Louis, MO, USA). Mouse monoclonal anti-MDM2 (SMP14), anti-p53 (DO-1) and anti-GAPDH (0411) antibodies were purchased from Santa Cruz Biotechnology, Inc. (Dallas, TX, USA). Rabbit polyclonal antibody against Lamin A/C (NB100-56649) was obtained from Novus Biologicals, LLC (Centennial, CO, USA). DyLight 800/680 conjugated (for western blots) and Alexa Fluor 488/555 tagged (for confocal micoscopy) secondary antibodies anti-mouse, anti-rabbit, anti-sheep IgG (H+L) were purchased from Thermo Fisher Scientific Inc. (Waltham, MA, USA).

### Cell lines and Cell culture

HEK293 cells were obtained from Rupak Dutta (Indian Institute of Science Education and Research, Kolkata, India). HEK293T cells were purchased from GE Healthcare Dharmacon Inc., Lafayette, CO, USA. Both HEK293 and HEK293T cells were maintained in Dulbecco’s modified Eagle’s medium (DMEM) (Gibco*/*Invitrogen, Inc., USA) supplemented with 10% fetal bovine serum (FBS) (Gibco*/*Invitrogen, Inc., USA), 1% Penicillin-Streptomycin Solution (Sigma-Aldrich Corp., St. Louis, MO, USA). EBV-positive Burkitt’s lymphoma (BL) cell line Namalwa was obtained from National Centre for Cell Science, (NCCS, Pune, Govt. of India). EBV-negative BL lines – DG75, BJAB and BJAB stably expressing EBNA3A and EBV positive BL line Raji cells were obtained from Elliott Kieff (Harvard Medical School, USA). *In vitro* EBV transformed LCL clones (LCL#1 and LCL#89), EBV-negative BL line BJAB, BJAB clones stably expressing either vector or EBNA3C cDNA (E3C#7) and lentivirus-mediated stable EBNA3C knockdown (LCL#Sh-E3C) or control (LCL#Sh-Con) lymphoblastoid cell lines (LCLs) were previously described. LCL#Sh-E3C and LCL#Sh-Con cells were freshly generated in LCL#1 as described below. BJAB-vector and BJAB-EBNA3C cells were maintained in complete RPMI media supplemented with 500 μg/ml G418 (Sigma-Aldrich Corp., St. Louis, MO, USA). LCL#sh-control and LCL#sh-EBNA3C cells were maintained in complete RPMI media supplemented with 1 μg/ml puromycin (Sigma-Aldrich Corp., St. Louis, MO, USA). LCLs were maintained in RPMI 1640 (Gibco*/*Invitrogen, Inc., USA) supplemented with 10% FBS and 1% Penicillin-Streptomycin Solution. Unless otherwise stated all above-mentioned cells were cultured at 37^0^C in a humidified environment supplemented with 5% CO_2_.

### Western blotting

Unless and otherwise stated, ∼10 x 10^6^ adherent or B-cells were harvested, washed with ice cold 1X PBS (Invitrogen, Thermo Fisher Scientific Inc., Waltham, MA, USA), and subsequently lysed in 0.5ml ice cold RIPA buffer (Sigma-Aldrich Corp., St. Louis, MO, USA) with protease inhibitor cocktail (Cell Signaling Technology Inc., Danvers, MA, USA). Protein concentrations were estimated by Bradford reagent (BIO-RAD, Hercules, CA, USA). Samples were boiled in 2 X laemmli buffer (BIO-RAD, Hercules, CA, USA), fractionated by SDS-PAGE and transferred to a 0.45 µm either nitrocellulose or low autofluorescence PVDF membranes (BIO-RAD, Hercules, CA, USA). The membranes were then probed with specific antibodies followed by incubation with appropriate infrared-tagged/DyLight secondary antibodies and viewed on an Odyssey CLx Imaging System (LiCor Inc., Lincoln, NE, USA). Image analysis and quantification measurements were performed using the Odyssey Infrared Imaging System application software (LiCor Inc., Lincoln, NE, USA).

### Transfection

Unless and otherwise stated, ∼10 x 10^6^ HEK293 cells were harvested, resuspended with 450 µL Opti-MEM medium (Invitrogen, Thermo Fisher Scientific Inc., Waltham, MA, USA), mixed with appropriate plasmids in a 0.4-cm gap cuvette (BIO-RAD, Hercules, CA, USA) and transfection was performed by electroporation using a Gene Pulser II electroporator (BIO-RAD, Hercules, CA, USA) at 210 V and 975 µF for Western Blot analyses. Cells were harvested 36 h post-transfection for western blot analyses, unless and otherwise stated. For confocal microscopy, cells were transfected with appropriate plasmids using Lipofectamine 3000 according to manufacturer’s protocol (Invitrogen, Thermo Fisher Scientific Inc., Waltham, MA, USA). 24 h post-transfection cells were subjected to live-cell confocal microscopy analyses.

### Stability assay

∼5 x 10^6^ EBV transformed B-lymphocytes (LCL#1) or EBNA3C stably expressing BJAB cells (BJAB-E3C#7) were treated with 100 µg/ml cyclohexamide (Sigma-Aldrich Corp. St. Louis, MO, USA) alone (with DMSO control) or in the presence of 1 µM MG132 (Abcam Inc, Cambridge, UK) or 50 µM Chloroquine (Sigma-Aldrich Corp. St. Louis, MO, USA) in normal growth medium for 24 h. Subsequently, lysates were prepared in 1 x RIPA buffer at indicated time periods and subjected to western blot analyses with appropriate antibodies. Band intensities were quantified by Odyssey imager (LiCor Inc., Lincoln, NE, USA).

### Confocal Microscopy

∼1 x 10^6^ HEK293 cells grown at 70% confluency on 35 mm dish embedded with 12 mm glass viewing area (Nunc, Thermo Fisher Scientific Inc., Waltham, MA, USA) were transfected with appropriate plasmids using Lipofectamine 3000. 24 h post-transfection cells were either left untreated (DMSO control) or treated with 20 µM MG132 for another 4-6 h and subsequently subjected for live-cell confocal analyses. For live-cell imaging, Hoechst 3342 solution (Thermo Fisher Scientific Inc., Waltham, MA, USA) and LysoTracker Red DND-99 (Invitrogen, Thermo Fisher Scientific Inc., Waltham, MA, USA) were used to stain the nucleus and the acidic lysosomes, respectively. For LCLs, approximately 1 million cells either left untreated (DMSO control) or treated with 1 µM MG132 for 12 h were harvested in microcentrifuge tubes, washed with 1 X PBS, fixed and permeabilized with 4% paraformaldehyde (Sigma-Aldrich Corp. St. Louis, MO) and 0.1% Triton X-100 (Sigma-Aldrich Corp. St. Louis, MO, USA) followed by blocking with 5% BSA (BIO-RAD, Hercules, CA) at room temperature for 1 h. Cells were further incubated with appropriate primary antibodies for overnight at 4^0^C. Nucleus was counterstained with 4’, 6’,-diamidino-2-phenylindole (DAPI; BIO-RAD, Hercules, CA, USA) for 30’ at room temperature. Alexa Fluor labeled secondary antibodies diluted in blocking buffer (1:2000) were incubated at room temperature for another 1 h. Cells were then washed for three times with 1 X PBS and mounted using an antifade mounting media (Sigma-Aldrich Corp. St. Louis, MO, USA) in 8-well chamber slides (Thermo Fisher Scientific Inc., Waltham, MA, USA) for confocal microscopy. The images were obtained by a Leica DMi8 Confocal Laser Scanning Microscope and analyzed in the Leica Application Suite X (LAS X) software (Leica Microsystems GmbH, Wetzlar, Germany). The intensities of stains were sometimes also analyzed by ImageJ software (https://imagej.nih.gov/ij/).

### Real-Time quantitative PCR (qPCR)

Total RNA was isolated from approximately 10 million LCLs (both LCL#1 and LCL#89) either left untreated (DMSO control) or treated with 1 µM MG132 for 12 h using TRIzol reagent according to the manufacturer’s instructions (Invitrogen, Thermo Fisher Scientific Inc., Waltham, MA, USA), followed by cDNA preparation using iScript cDNA synthesis kit (BIO-RAD, Hercules, CA, USA) as per manufacturer’s protocol. RNA and cDNA quality and quantity were checked using Synergy™ H1 Multimode Microplate Reader (BioTek Instruments, Inc., VT, USA). qPCR analysis was performed using iTaq Universal SYBR Green Supermix (BIO-RAD, Hercules, CA, USA) in CFX Connect™ Real-Time PCR detection System (BIO-RAD, Hercules, CA, USA) with the following thermal profile – 1 cycle: 95^0^C for 10 min; 40 cycles: 95^0^C for 15 sec followed by 60^0^C for 1 min; and finally the dissociation curve at – 95°C for 1 min, 55°C 30 min, and 95°C for 30 sec. Unless and otherwise stated, each sample was performed in duplicate and calculation was made using a 2^−ΔΔCT^ method to quantify relative expression compared with housekeeping gene control. The real time PCR primers used in this study were designed from Primer-BLAST (https://www.ncbi.nlm.nih.gov/tools/primer-blast/) and listed in Table S2. Real-time PCR primers were obtained from Integrated DNA Technologies, Inc. (Coralville, IA, USA).

### RNA-Sequencing and data analyses

Approximately 1 µg of total RNA isolated from LCLs (both LCL#1 and LCL#89) either left untreated (DMSO control) or treated with 1 µM MG132 for 12 h was subjected to whole transcriptome enrichment through selective depletion of 99.9% of ribosomal RNA using RiboMinus™ Eukaryote Kit for RNA-Seq (Thermo Fisher Scientific Inc., Waltham, MA, USA) followed by library generation using Ion Total RNA-Seq Kit v2 (Thermo Fisher Scientific Inc., Waltham, MA, USA). Transcriptome analysis was performed in Ion S5^TM^ XL System (Thermo Fisher Scientific Inc., Waltham, MA, USA). Reads generated from Ion S5 XL were subjected to further analysis with a minimum read length of 25 bp. The filtered in reads were aligned to human genome (hg19) using two-step alignment method. Reads were aligned using STAR and Bowtie software using default parameters. Both Alignment data were merged together using Picard. Reads aligning to reference genome were provided in bam format. Using the encode gene annotations for human genome (hg19), aligned reads were counted for each gene and were represented in standard format for all 4 samples. RNASeqAnalysis plugin (v5.2.0.5) was utilized to perform the analysis and produce gene counts for all the samples.

Using DESeq2 package from R, study for differential expression analysis was performed between sample conditions. It followed the negative binomial distribution for studying gene counts and tested each gene from samples for expression. Genes were selected based on p-value as <=0.05 and log_2_ Fold Change as 2 and above (up-regulated) and −2 and below (down-regulated). Differentially expressed gene sets were segregate together and subsequently uploaded on DAVID (<u>D</u>atabase for <u>A</u>nnotation, <u>V</u>isualization and Integrated <u>D</u>iscovery, version 6.8) webserver (https://david.ncifcrf.gov/) for further analyses. Features observed in different databases are clustered together for functional analysis. Gene Ontology (GO) was selected from the hits table for DAVID clustering. The clusters were represented through statistical analysis providing P-value and FDR (false discovery rate) in order to select functional significant clusters.

### Induction of EBV lytic cycle from LCLs

∼10 x 10^6^ LCLs (both LCL#1 and LCL#89) were induced to release virus in complete RPMI 1640 medium containing 20 ng/ml TPA (Sigma-Aldrich Corp. St. Louis, MO, USA) and 3 mM sodium butyrate (Sigma-Aldrich Corp. St. Louis, MO, USA) for 24 h. To understand the role of proteasomal inhibition in EBV lytic cycle induction, LCLs were either left untreated (DMSO control) or treated with 1 µM MG132 for 24 h. Cells were then harvested and subjected for genomic DNA isolation using “Blood & Cell Culture DNA Mini Kit” (Qiagen, Hilden, Germany). Reactivation of LCLs was determined by real-time quantitative PCR (qPCR) analyses of EBV DNA targeting *Bam*HI W genomic portion (Forward primer: 5’-CCCAACACTCCACCACACC-3’; reverse primer: 5’-TCTTAGGAGCTGTCCGAGGG-3’). As described above, the primers were synthesized from Integrated DNA technologies, Inc. (Iowa, USA).

### Generation of Beclin 1 and EBNA3C knockdown cells

∼10 x 10^6^ HEK293 cells were transfected with pTripZ vector containing sh-Beclin1 sequence under doxycycline inducible promoter by electroporation. Sh-RNA expression was initiated using 1 µg/ml doxycycline (Sigma-Aldrich Corp. St. Louis, MO, USA) and transfected cells were selected using 2 µg/ml of puromycin (Sigma-Aldrich Corp. St. Louis, MO, USA) for 7 days. Without doxycycline treatment was served as control.

For EBNA3C knockdown in LCLs, ∼10 x 10^6^ HEK293T cells (70% confluency) were first transfected with lentiviral packaging vectors pCMV-VSV-G, pRSV-Rev and pMDLg/pRRE along with pGipz vector either expressing control sh-RNA (5′-TCTCGCTTGGGCGAGAGTAAG-3′) or sh-EBNA3C (5’-CCATATACCGCAAGGAAT-3’). Lentivirus production and transduction LCLs were essentially carried out as previously described. 12 h post-transfection, the medium was changed with fresh DMEM supplemented with 10 mM sodium butyrate for virus induction for 8 h. Lentivirus was collected twice per day (every 10-12 h) for 2 days in RPMI 1640 supplemented with 10% FBS and 10 mM HEPES, filtered through 0.45 μM filters (Corning Inc., NY, USA) to avoid any cell contamination. The supernatant containing virus was concentrated (100x) by ultra-centrifugation at 70,000 × g for 2.5 h and subsequently used for infection in the presence of 10 μg/ml Polybrene solution (Sigma-Aldrich Corp. St. Louis, MO, USA). 3-days post-infection cells were selected using 1 μg/ml puromycin for 14 days. Selected cells were grown in complete RPMI supplemented with 1 μg/ml puromycin for 1 month before subjected to western blot analyses and cell viability assays.

### Colony formation assays

∼10 x 10^6^ HEK293 stably transfected with pTripz-Sh-Beclin1 were either left untreated or treated with doxycycline to express the sh-RNA. 72 h post-treatment approximately 1 x 10^6^ cells were further treated with DMSO control or 20 µM MG132 for 4 h in 100 mm petri-plates and selected in complete DMEM supplemented with 2 µg/ml puromycin. Two weeks later, cells were fixed with 4% paraformaldehyde paraformaldehyde (Sigma-Aldrich Corp. St. Louis, MO) at room temperature for 0.5 h and stained with 0.1% crystal violet solution (Sigma-Aldrich Corp. St. Louis, MO, USA) for 0.5 h. The plates were scanned using Odyssey CLx Imaging System (LiCor Inc., Lincoln, NE, USA) and the relative colony number was measured by Image J software.

### Co-immunoprecipitation assay

∼15 x 10^6^ HEK293 cells transiently transfected with the indicated expression plasmids or 20 million LCLs were harvested, washed twice with ice cold 1 x PBS and subsequently lysed with 0.5 ml ice cold RIPA buffer, supplemented with protease inhibitors for 1 h on ice with brief vortexing every 5-10 minutes. The lysates were centrifuged at 13,000 rpm for 15 min at 4°C. 5 μl of each sample was taken out separately for measuring protein concentration using Bradford protein assay reagent (BIO-RAD, Hercules, CA, USA). 5% of the lysates were boiled with 2 x laemmli buffer (BIO-RAD, Hercules, CA, USA) and used as input. Lysates were precleared with control antibody plus 30 µL of 1:1 Protein-A/G magnetic beads (BIO-RAD, Hercules, CA, USA) for 1 h at 4^0^C. The protein of interest was captured by rotating the remaining lysate with 1 µg of specific antibody overnight at 4^0^C. Immuno-complexes were captured with 30 μL magnetic Protein-A/G beads, pelleted and washed 5 x with ice cold RIPA buffer. Input lysates and IP complexes were boiled in 2 x laemmli buffer, fractionated by SDS-PAGE and transferred to either 0.45 µm nitrocellulose or PVDF membrane for western blot analyses as described above on an Odyssey imager.

### *In vivo* ubiquitination assay

∼15 x 10^6^ HEK293 cells were transfected with flag-tagged EBNA3C constructs expressing either full-length protein (residues 1-992) or various domains N-terminal (residues 1-365), middle part (residues 366-620) and the C-terminal (residues 621-992). 36 h post-transfection cells were further incubated with 20 µM MG132 for an additional 4 h before harvesting. After washing with 1 x PBS, cells were lysed in 100 µl RIPA buffer containing 1% SDS, boiled for 10 minutes to disrupt any protein-protein interaction. Each lysate was subsequently mixed with 900 µl of RIPA buffer without SDS to dilute the SDS concentration to 0.1%, incubated at 4^0^C on rotator for 1 h and centrifuged at 13,000 rpm for 15 min at 4°C. The supernatant was collected to a new tube and precleared with control antibody plus 30 µL of 1:1 Protein-A/G magnetic beads for 1 h at 4^0^C. Flag-tagged EBNA3C proteins were immunoprecipitated with 2 µg M2 antibody for overnight at 4^0^C, captured by 1:1 Protein-A/G magnetic beads, washed 5 x with ice cold RIPA buffer, and boiled in 2 x laemmli buffer. The extent of ubiquitination of flag-tagged proteins was determined by SDS-PAGE followed by western blot analysis using indicated antibodies against total ubiquitin or specific to K48- or K63-linked polyubiquitination.

### Subcellular fractionation assay

**∼**10 x 10^6^ HEK293 cells transiently transfected for 36 h with the indicated EBNA3C expression plasmids were either left untreated (DMSO control) or treated with 20 µM MG132. Cells were harvested and subsequently subjected to subcellular fractionation as per manufacturer’s protocol (BIO-RAD, Hercules, CA, USA). Nuclear and cytoplasmic protein fractions were measured by Bradford protein assay (BIO-RAD, Hercules, CA) and ∼50 µg of total protein was resolved by SDS-PAGE. The efficiency of nuclear and cytoplasmic fractionation was confirmed by western blot analyses against nuclear protein Histone or lamin A/C and cytoplasmic protein GAPDH, respectively.

### Soft agar assay

Soft agar assays were performed for LCLs (both LCL#1 and LCL#89). 0.75% agar (Sigma-Aldrich Corp. St. Louis, MO, USA) in 1 ml complete RPMI 1640 medium was poured into each well of 6-well plates (Corning Inc., NY, USA) and set aside to solidify. Next, 1 × 10^5^ LCLs either left untreated (DMSO control) or treated with MG132 (1-5 µM) or bortezomib (0.5-1 µM) for 24 h, were harvested, mixed with 1 ml complete RPMI 1640 medium supplemented with 0.36% agar and poured on the top of the hard agar (1%) layer. Two weeks later, colonies were stained with 0.1% crystal violet for 30 minutes and scanned using Odyssey CLx Imaging System (LiCor Inc., Lincoln, NE, USA) and the relative colony number was measured by Image J software. Prior to staining, each well was also photographed (bright-field) using a ZOE™ Fluorescent Cell Imager (BIO-RAD, Hercules, CA, USA).

### Cell viability assay

∼0.5 × 10^5^ cells plated into each well of the six-well plates (Corning Inc., NY, USA) were treated with either MG132 or bortezomib at increasing concentrations (0-10 µM) for the indicated time points at 37^0^C in a humidified CO_2_ chamber. Viable cells from each well were measured by Trypan blue exclusion method using an automated cell counter (BIO-RAD, Hercules, CA). Experiments were performed in duplicate and were independently repeated two times.

### Measurement of intracellular ROS

∼1 × 10^5^ LCLs (both LCL#1 and LCL#89) either left untreated (DMSO control) or treated with 0.5 µM MG132 in the presence and absence of ROS (<u>R</u>eactive <u>O</u>xygen <u>S</u>pecies) scavenger 1 mM N-Acetyl-L-cysteine (NAC) for 24 h were harvested, suspended in PBS in a 96-well plate (Corning Inc., NY, USA). The fluorescent probe DCFH-DA (Sigma-Aldrich Corp. St. Louis, MO, USA) was used to detect intracellular ROS levels. After incubation with 20 μM DCFH-DA for 30 min at 37^0^C and the fluorescence was measured by Synergy™ H1 Multimode Microplate Reader (BioTek Instruments, Inc., VT, USA) using the blue filter (485 nm) for excitation and the green filter (528 nm) for emission. Experiments were performed in triplicate and were independently repeated two times.

### Statistical analysis

All the data represented are as the mean values with standard deviation (SD). Statistical significance of differences in the mean values was analyzed using the Student’s t-test two tail distribution and equal variances between two samples. P-value below 0.05 was considered as significant (*P < 0.05; **P < 0.01; ***P < 0.001; NS, not significant).

## Acknowledgements

We sincerely thank to Elliott Kieff (Harvard Medical School, USA), Martin Rowe (University of Birmingham, UK), Jayanta Debnath (University of California, San Francisco, USA), Edward M. Campbell (Loyola University Chicago, USA), Robert A. Weinberg (Whitehead Institute for Biomedical research, Cambridge, USA), Rupak Dutta (Indian Institute of Science Education and Research, Kolkata, India), Nabendu Biswas (Presidency University, Kolkata, India) and National Centre for Cell Science (NCCS), Dept. of Biotechnology (DBT), Govt. of India for providing reagents, plasmids, cell lines and hybridomas. We thank Rupak Dutta (Indian Institute of Science Education and Research, Kolkata, India) and Shubhra Majumder (Presidency University, Kolkata, India) for careful review of the manuscript. We also thank Pralay Majumder (Presidency University, Kolkata, India) for technical support in using Microscopy Facility at Dept. Life Sciences, Presidency University, Kolkata, India. E.S.R. is a scholar of the Leukemia and Lymphoma Society of America. A.S. and B.B.D. are Wellcome Trust/DBT India Alliance Intermediate Fellows. C.B. and S.M. are the recipients of UGC-NET Senior and Junior Research Fellowship, respectively, India. S.B. is recipient of DST-Inspire Senior Research Fellowship and A.G. is recipient of CSIR-NET Senior Research Fellowship.

## Supporting Information

**Figure S1: Proteasome inhibitors induce degradation of EBV oncoproteins - EBNA3A and EBNA3C.** ∼10 x 10^6^ (A) BJAB stably expressing EBNA3A (BJAB#E3A), (B) BJAB stably expressing EBNA3C (BJAB#E3C), (C) LCL#89 cells either left untreated (DMSO control) or treated with (A-B) 1 µM MG132, (C) 0.5 µM bortezomib for 12 h, were harvested and subjected for western blot analyses using the indicated antibodies. GAPDH blot was used as loading control. Protein bands were quantified by Odyssey imager software and indicated as bar diagrams at the bottom of corresponding lanes.

**Figure S2: EBNA3C fractionation in the absence and presence of leptomycin B.** HEK293 cells transiently transfected with flag-tagged EBNA3C construct either left untreated (DMSO control) or treated with leptomycin B (LMB; 20 ng/ml) for 24 h, were subjected to subcellular fractionation as described in the “Materials and Methods” section. Protein bands were quantified by Odyssey imager software and indicated as bar diagrams at the bottom of corresponding lanes.

**Figure S3: The N-terminal domain of EBNA3C plays crucial role in forming complex with p62 and LC3B.** (A) HEK293 cells transiently transfected with expression plasmids for flag-tagged EBNA3C truncations (residues 1-365, 366-621 and 621-992) were subjected to co-immunoprecipitation study with anti-flag antibody. Western blots were performed with the indicated antibodies by stripping and reprobing the same membrane. * indicates IgG bands. (B) Schematic representation of EBNA3C truncations used in the co-immunoprecipitation experiment. (C-D) Summary and cartoon representation of the interaction study of different EBNA3C domains with p62 and LC3B.

**Figure S4: Caspase inhibitor does not rescue MG132-induced EBNA3C’s degradation.** LCLs were either left untreated (DMSO control) or treated with 1 µM MG132, 50 µM pan-caspase inhibitor Z-VAD(OMe)-FMK (Z-VAD) or MG132 plus Z-VAD. 24 h post-treatment cells were subjected for either (A) western blot analysis or (B) immunostaining with the indicated antibodies. Each panel in (B) is representative picture of two independent experiments and nuclei were counterstained by DAPI before mounting the cells. Scale bars, 5 µm. In (A) GAPDH blot was used as loading control and protein bands were quantified by Odyssey imager software and represented as bar diagrams at the bottom of corresponding lanes.

**Figure S5: MG132 mediated ROS generation has no role in EBNA3C degradation.** LCLs were either left untreated (DMSO control) or treated with 0.5 µM MG132, 1 mM N-Acetyl-L-cysteine (NAC) or MG132 plus NAC. 24 h post-treatment, cells were subjected for (A-B) ROS (<u>R</u>eactive <u>O</u>xygen <u>S</u>pecies) measurement using DCFH-DA fluorescent probe, (C) western blot analysis and (D) immunostaining with the indicated antibodies. (C) GAPDH blot was used as loading control and protein bands were quantified by Odyssey imager software and represented as bar diagrams at the bottom of corresponding lanes. Each panel in (D) corresponds to single experiment of two independent experiments and nuclei were counterstained by DAPI before mounting the cells. Scale bars, 5 µm.

**Figure S6: EBNA3C expression in different Cell lines.** ∼10 x 10^6^ cells were harvested, lysed in RIPA buffer and subjected to western blot analyses with anti-GAPDH (as loading control) and anti-EBNA3C antibodies.

**Table S1. RNA-Seq and Gene Ontology term analyses of DMSO and MG132 treated LCLs.** ∼10 x 10^6^ LCL#1 and LCL#89 either left untreated or treated with 1 µM MG132 for 12 h, were subjected to RNA-Seq followed by Gene Ontology (GO)-Term Enrichment analyses as described in the “Materials and Methods” section.

**Table S2. Real-time PCR primers.** Real-time PCR primers for both viral and cellular genes were designed from Primer-BLAST (https://www.ncbi.nlm.nih.gov/tools/primer-blast/). Reference Sequence ID (RefSeq ID) of each gene is mentioned. All primers were selected at annealing temperature of ∼60^0^C. RefSeq ID of each gene is mentioned.

